# Peripheral T-cell co-signalling states mark vulnerability and resilience to cerebral Aβ pathology

**DOI:** 10.64898/2026.08.03.742616

**Authors:** Anna Mallone, Dario Bachmann, Chiara Rickenbach, Maik Krüger, Tunahan Kirabali, Henrik Zetterberg, Maria Teresa Ferretti, Luka Kulic, Christoph Hock, Roger M. Nitsch, Federica Sallusto, Anton Gietl, Valerie Treyer, Christoph Gericke

## Abstract

Adaptive immune responses may influence vulnerability and resilience in Alzheimer’s disease (AD), but relevant T-cell states remain unclear. We profiled peripheral immune cells by mass cytometry in 200 participants across distinct age groups, early AD and exceptional old age without dementia, relating immune features to amyloid-β (Aβ) PET, plasma biomarkers and longitudinal structural and cognitive outcomes. Inducible T-cell co-stimulator (ICOS) expression across CD4 and CD8 memory T-cells was associated with cerebral Aβ pathology. In mild cognitive impairment (MCI), higher ICOS expression on CD8 memory T-cells strengthened the association between Aβ load and hippocampal atrophy. Aβ-derived peptides induced proliferative ICOS+CD25+ memory CD4 and CD8 T-cell responses predominantly in Aβ-positive participants in an independent cohort. Conversely, higher programmed cell death protein 1 (PD-1) expression on CD8 effector-memory T-cells was associated with attenuated Aβ-related episodic-memory decline in exceptionally old participants and was higher in stable MCI than in MCI-to-AD converters. These findings identify distinct co-stimulatory and co-inhibitory T-cell correlates of vulnerability and resilience.

## Introduction

Alzheimer’s disease (AD) is neuropathologically defined by amyloid-β (Aβ) plaques and tau neurofibrillary tangles, accompanied by broader cellular and molecular processes that may ultimately result in neurodegeneration and cognitive decline.^1,2^ Aβ and tau pathologies begin years before cognitive symptoms^3,4^ and can be detected using positron emission tomography (PET) and cerebrospinal fluid (CSF) biomarkers,^5^ while plasma Aβ and phosphorylated tau (p-tau) species provide accessible blood-based measures.^6^ Biomarker-positive cognitively unimpaired individuals are at increased risk of progression to mild cognitive impairment (MCI) or dementia, particularly when both Aβ and tau pathology are present,^7^ yet progression to AD dementia is not inevitable.^8^ The prevalence of cerebral Aβ pathology and the incidence of dementia increase with age,^9,10^ but substantial AD pathology can coexist with preserved cognition,^11,12^ even among individuals at exceptional old age.^13,14^ This dissociation between pathological burden and cognitive impairment highlights resilience mechanisms that modify the clinical expression of AD pathology.^5,15^

Resilience to AD pathology is likely multifactorial. Cognitive reserve and lifetime experience,^16^ physical activity and lifestyle,^16,17^ sleep,^18^ sex,^17,19^ vascular health^13,20^ and genetic background, including *APOE*,^13,21^ have all been associated with differences in pathology burden or cognitive outcome. Emerging evidence also places immune factors among these potential modifiers. Genetic analyses have implicated immune-response and immune-system activation pathways in resilience to AD and cerebral Aβ pathology.^22,23^ Human single-cell studies have identified distinct microglial transitions associated with resilience at the inflection point between Aβ- and tau-associated pathological processes^24^ and peripheral immune signatures associated with exceptional longevity.^25^

Immune mechanisms are increasingly recognized as integral components of AD pathobiology.^2^ Current frameworks incorporate interactions among resident brain cells, brain-border compartments and peripheral immune populations, including adaptive immune cells.^26,27^ T-cells have been observed in human AD brains for decades.^28^ Later studies identified predominantly CD8 extravascular T-cells in AD hippocampi associated with tau rather than Aβ pathology^29^ and clonally expanded CD8 T-cells in AD CSF.^30^ T-cell infiltration has also been reported in mouse models of cerebral Aβ pathology.^31^ In blood, alterations in CD8 effector-memory T-cells re-expressing CD45RA (TEMRA) have been detected in cognitively unimpaired and MCI individuals with cerebral Aβ accumulation.^32,33^ Experimental models of Aβ and tau pathology have assigned divergent functions to CD8 T-cells.^34–37^ Together, these findings shift the central question from whether T-cells play a role in AD pathobiology to which subsets and states are involved, how they act across disease stages and whether their effects are beneficial or detrimental. These questions cannot be resolved from cell abundance alone.

Antigen specificity and co-signalling provide complementary routes to resolve this functional heterogeneity. Human studies have detected T-cell reactivity to Aβ-derived peptides in older individuals and patients with AD,^38,39^ and our previous work identified Aβ peptide-responsive memory CD4 T-cells through induction of the co-stimulatory receptors OX40 and 4-1BB.^33^ These findings established activation-induced co-stimulatory receptor expression as a means of identifying Aβ-responsive T-cells, but left open whether broad immune profiling would reveal additional co-signalling phenotypes associated with Aβ pathology and subsequent structural or cognitive outcomes. Among co-signalling receptors, inducible T-cell co-stimulator (ICOS) is a CD28-family co-stimulatory receptor that supports T-cell proliferation, survival and differentiation,^40,41^ whereas programmed cell death protein 1 (PD-1) is a co-inhibitory receptor that constrains TCR and CD28 signalling following ligand engagement.^41,42^ How T-cell phenotypes marked by these receptors relate to cerebral Aβ pathology, neurodegenerative vulnerability or resilience has not been established in detail.

Here, we used high-dimensional mass cytometry to broadly profile peripheral immune cells from cerebral Aβ-stratified cognitively unimpaired, MCI and oldest-old participants aged ≥85 years without dementia. We related immune profiles to plasma and imaging biomarkers, and to longitudinal structural and cognitive outcomes. Aβ peptide-induced T-cell responses were additionally examined in an independent study population. We identify an ICOS-centred activation state across CD4 and CD8 memory T-cells associated with cerebral Aβ pathology and, in MCI, greater Aβ-associated hippocampal atrophy. By contrast, higher PD-1 expression on CD8 effector-memory T-cells was associated with attenuated Aβ-related episodic-memory decline in oldest-old participants and distinguished longitudinally stable MCI participants from MCI-to-AD converters. These findings identify co-stimulatory and co-inhibitory T-cell states as separable correlates of vulnerability and resilience.

## Methods

### Study design and participants

#### Study population 1

Study population 1 comprised peripheral blood mononuclear cells (PBMCs), plasma and serum from 200 participants assigned to four groups: young adults (Y; n = 10), cognitively unimpaired older adults (CU; n = 95), participants with mild cognitive impairment (MCI; n = 47) and oldest-old participants without dementia (OO; n = 48) (**Table 1 and Extended Data Table 1**). Anonymous blood samples from healthy adults younger than 40 years were obtained from the local blood donation service in Schlieren, Zurich, Switzerland. Only age and immune-cell profiling data were available for these participants. The Y group was therefore included exclusively as a reference population for the analysis of age-associated immune-cell differences and did not undergo clinical, neuropsychological, imaging, serological or plasma biomarker assessments. CU and MCI participants were enrolled in the monocentric ID-Cog study, an ongoing prospective, non-clinical cohort at the Center for Prevention and Dementia Therapy, University of Zurich. CU participants were aged >50 years and were classified as cognitively unimpaired based on comprehensive neuropsychological assessment. MCI was diagnosed according to established consensus criteria.^43^ Participants with severe immune-modifying disorders, including current or recent cancer and Hashimoto’s disease, were excluded. Detailed inclusion and exclusion criteria and cohort characteristics have been reported previously.^33,44,45^ The OO cohort comprised volunteers aged ≥85 years who had no dementia at baseline following a comprehensive clinical assessment and showed preserved everyday functioning. Because reliable normative ranges for neuropsychological performance are limited in this age range, OO participants were not further classified as CU or MCI. Detailed inclusion and exclusion criteria for the OO cohort have been published previously.^16^ Diagnostic assessments were performed independently of, and before, the immune-cell analyses.

**Table 1:** Demographics and clinical characteristics of study population 1.

| Variable | Y | CU– | CU+ | MCI– | MCI+ | OO– | OO+ |
| --- | --- | --- | --- | --- | --- | --- | --- |
| Number of subjects, n | 10 | 50 | 45 | 26 | 21 | 20 | 28 |
| Baseline age, years, mean ± SD | 31.80 ± 4.02 | 67.42 ± 6.35 | 68.29 ± 6.57 | 70.81 ± 9.04 | 74.64 ± 7.28 | 88.86 ± 3.35 | 88.34 ± 2.82 |
| Sex, n male/female | — | 23/27 | 35/10 | 17/9 | 11/10 | 13/7 | 20/8 |
| APOE ε4 carriers, % of group | — | 20.00% | 37.78% | 3.85% | 33.33% | 10.00% | 14.29% |
| **Neuropsychology / Cognitive testing** |  |  |  |  |  |  |  |
| Baseline MMSE, median [IQR] | — | 30.00 [1.00] | 29.00 [1.00] | 28.50 [1.00] | 28.00 [2.00] | 29.00 [1.50] | 29.00 [1.00] |
| Baseline episodic memory performance, z-score, median [IQR] | — | 0.12 [0.75] | -0.04 [1.15] | -0.85 [0.93] | -1.39 [2.40] | -1.00 [1.47] | -1.23 [1.63] |
| Years of NP follow-up, mean ± SD | — | 5.23 ± 1.84 | 6.08 ± 1.82 | 5.26 ± 1.55 | 5.01 ± 2.61 | 3.31 ± 2.68 | 2.76 ± 2.24 |
| **Baseline AD pathology biomarkers** |  |  |  |  |  |  |  |
| Aβ PET FMM Centiloid, median [IQR] | — | 1.23 [8.57] | 19.98 [18.34] | 0.66 [11.15] | 38.92 [58.49] | 2.56 [12.57] | 29.76 [51.85] |
| Plasma Aβ42/Aβ40, median [IQR] | — | 0.068 [0.013] | 0.062 [0.023] | 0.071 [0.015] | 0.059 [0.011] | 0.063 [0.015] | 0.054 [0.015] |
| Plasma p-tau217, pg/ml, median [IQR] | — | 1.25 [1.30] | 1.34 [1.66] | 1.35 [1.14] | 2.95 [2.12] | 1.69 [1.31] | 2.67 [2.01] |

CU, MCI and OO participants underwent cerebral Aβ PET imaging using [18F]-flutemetamol tracer. Serum was used for measurement of antiviral antibody titres and C-reactive protein (CRP), whereas plasma was used for measurement of AD-related biomarkers. Additional assessments included apolipoprotein E (*APOE)* genotyping, MRI and longitudinal neuropsychological testing. Analysis-specific sample sizes varied according to sample and data availability and are reported in the figures, source data and **Extended Data Table 1**.

No statistical method was used to predetermine the overall sample size. Minimum sample size for the CU population in the context of adaptive immune changes was estimated previously.^46^ Beyond this, all eligible samples with sufficient material and available quality-controlled data were included. Samples were randomized during processing before live-cell barcoding and pooling. The 190 CU, MCI and OO samples were processed within one barcoded mass-cytometry experiment. The Y samples were acquired subsequently and harmonized computationally with the principal dataset.^47^

#### Study population 2

Study population 2 was derived from an independent previously established cohort and was used as a validation sample set using the *in vitro* antigen-stimulation workflow. The population comprised cognitively unimpaired control participants (CU; n = 14) and participants with MCI (n = 10), diagnosed according to consensus criteria.^43^ Participants with severe immune-modifying disorders, including current or recent cancer and Hashimoto’s disease, were excluded. Cryopreserved PBMCs and plasma were available for the present analyses. Participants underwent cerebral Aβ PET imaging using [11C]-Pittsburgh compound B (PiB) tracer, plasma AD biomarker measurements, *APOE* genotyping and routine assessment of CRP. Numbers of participants contributing to individual stimulation analyses are reported in the corresponding results and source data (**Table 2 and Extended Data Table 2**).

**Table 2:** Demographics and clinical characteristics of study population 2.

| Variable | CU– | CU+ | MCI– | MCI+ |
| --- | --- | --- | --- | --- |
| Number of subjects, n | 4 | 10 | 4 | 6 |
| Baseline age, years, mean ± SD | 73.00 ± 3.65 | 76.70 ± 3.71 | 71.00 ± 3.74 | 75.83 ± 4.96 |
| Sex, n male/female | 2/2 | 5/5 | 3/1 | 4/2 |
| APOE ε4 carriers, % of group | 0% | 55.56% | 50.00% | 83.33% |
| **Neuropsychology / Cognitive testing** |  |  |  |  |
| Baseline MMSE, median [IQR] | 29.00 [0.25] | 30.00 [1.00] | 28.50 [1.50] | 27.50 [3.75] |
| **Baseline AD pathology biomarkers** |  |  |  |  |
| Aβ PET PiB Centiloid, median [IQR] | -1.74 [2.65] | 52.83 [27.68] | -3.47 [7.79] | 88.68 [25.42] |
| Plasma Aβ42/Aβ40, median [IQR] | 0.055 [0.005] | 0.045 [0.005] | 0.045 [0.016] | 0.040 [0.009] |
| Plasma p-tau181, pg/ml, median [IQR] | 1.29 [0.52] | 1.96 [0.51] | 1.55 [0.25] | 2.73 [2.12] |

### Longitudinal diagnostic classification

Longitudinal diagnostic trajectories were constructed using the diagnosis recorded at baseline and all available valid follow-up diagnoses. Diagnostic categories were CU, MCI, probable AD dementia late-onset (PRADL) and OO without dementia. PRADL was diagnosed according to NINCDS-ADRDA criteria and ICD-10 criteria.^48,49^

Participants were required to have at least one valid follow-up diagnosis to be classified as longitudinally stable or as converters. Stable CU, MCI and OO participants retained their respective baseline diagnosis at all valid follow-up assessments. Persistent conversion from CU to MCI was defined by the first MCI diagnosis followed by no subsequent return to the baseline diagnosis. Conversion from MCI or OO to PRADL required all subsequent valid diagnoses to remain PRADL. Diagnostic changes followed by a return to the baseline category were classified as transient or non-persistent. Participants with valid follow-up information who met neither the stable nor persistent-converter definitions were classified as other non-stable followed participants. Specific diagnostic transition counts and the resulting classification groups are reported in **Extended Data Table 3**.

### APOE genotyping

*APOE* genotyping was performed using a restriction-isotyping protocol^50^ or by Sanger sequencing (Microsynth AG, Switzerland).

### PET imaging and image analysis

#### [18F]-Flutemetamol Aβ-PET in study population 1

Aβ-PET images were acquired on a Signa PET/MR system (GE Healthcare). Participants received approximately 140 MBq [18F]-flutemetamol (Vizamyl, GE Healthcare), and PET data were acquired from 80 to 110 min after injection. A three-dimensional T1-weighted BRAVO image with an isotropic voxel size of 1 mm was acquired in parallel for anatomical segmentation. PET images were processed using PMOD NeuroTool version 4.0 (PMOD Technologies) according to standard Centiloid procedures.^51^ Reconstructed PET frames acquired from 85 to 105 min after injection were averaged. A global cortical Aβ standardized uptake value ratio (SUVR) was calculated from a composite region comprising left and right frontal, temporal and parietal cortices and the anterior and posterior cingulate cortices, normalized to uptake in cerebellar grey matter. SUVR values were converted to the Centiloid scale. Cerebral Aβ positivity was defined as Centiloid >12, a research threshold reflecting the transition from absence of detectable pathology to subtle Aβ deposition.^52,53^

#### [11C]-PiB Aβ-PET in study population 2

Aβ-PET images were acquired over 70 min on whole body PET/CT systems in 3D mode (Discovery RX and Discovery STE, GE Healthcare). Participants received approximately 350 MBq [11C]-PiB tracer. Anatomical 3D T1-weighted MRI was acquired on a 3T Philips Achieva system using a turbo field-echo sequence with a voxel size of approximately 0.94 × 0.94 × 1 mm. Images were processed using PMOD NeuroTool version 3.7. T1-weighted images were segmented and cortical regions were defined using the adult brain maximum-probability atlas implemented in PMOD. PET images were co-registered to the baseline MRI using the early 6-min PET frames, which approximate a perfusion image. Co-registration and regional delineations were visually inspected and corrected where required. Late-phase cortical uptake was calculated from frames acquired between 50 and 70 min after injection. A global cortical SUVR was obtained using the Centiloid cortical composite and cerebellar cortex as reference region and was subsequently converted to the Centiloid scale. Aβ positivity was defined as Centiloid >12. ^52,53^

### MR imaging and image analysis

#### T1-weighted MRI and hippocampal measurements

T1-weighted MRI scans were acquired on 3T GE Discovery MR750w or GE SIGNA Premier systems using 32- or 48-channel head coils. High-resolution 3D fast spoiled-gradient recalled images were acquired with an isotropic voxel size of 0.5 mm, sagittal orientation, repetition time of 11 ms, echo time of 5.2 ms, inversion time of 600 ms and flip angle of 8°. Images underwent quality assessment using the CAT12 toolbox,^54^ which was also used to derive total intracranial volume (TIV). Baseline hippocampal volume was segmented using the automatic segmentation of hippocampal subfields software.^55^ Hippocampal volume was expressed as a percentage of baseline TIV. Values were previously reported for the whole cohort.^19^ Longitudinal hippocampal change was estimated using pairwise longitudinal registration of baseline and follow-up T1-weighted images in SPM12. This procedure generated a midpoint-average image and a 3D Jacobian rate map reflecting the annualized rate of volumetric change. Hippocampal masks generated using the automatic segmentation of hippocampal subfields were applied to the Jacobian rate maps, and the median annualized rate across hippocampal voxels was used as the longitudinal hippocampal outcome.

#### FLAIR MRI and WMH measurements

Fluid-attenuated inversion recovery (FLAIR) images were acquired in parallel with Aβ-PET on a 3T Signa PET/MR scanner (GE Healthcare). High-resolution 3D cube FLAIR images were acquired with a voxel size of 0.48 × 0.48 × 0.6 mm, repetition time of 6,502 ms, echo time of 130.7 ms, inversion time of 1,962 ms and flip angle of 90°.

White-matter hyperintensities (WMHs) were segmented using the deep-learning-based MIAC Automated Region Segmentation pipeline^56^ employing the nnU-Net algorithm.^57^ All lesion masks were visually inspected. Global WMH volume was expressed as a percentage of baseline TIV. Longitudinal global WMH change was calculated as follow-up minus baseline WMH volume divided by the follow-up duration.

### Neuropsychological assessment

Details of the neuropsychological assessment were published previously.^17,19,32^ Global cognitive performance was characterized using the Mini-Mental State Examination (MMSE). Cognitive performance was assessed approximately annually for up to 8.78 years. The principal longitudinal cognitive outcome was an episodic-memory composite derived from immediate recall, delayed recall of words and figures, and recognition measures from the Consortium to Establish a Registry for Alzheimer’s Disease (CERAD) neuropsychological battery. Individual test scores were standardized using the baseline mean and standard deviation of CU participants, the same reference values were applied at all longitudinal time points. Individual rates of longitudinal episodic memory change (= ‘cognitive change’) were estimated in the combined longitudinal ID-Cog and OO datasets, including participants beyond study population 1, using a linear mixed-effects model (LMM) fitted by restricted maximum likelihood (REML) with the *nlme* package in R. The model included years from baseline as the sole fixed effect, with participant-specific random intercepts and random slopes. Participant-specific episodic memory slopes were obtained by adding the fixed population-level time coefficient to each participant’s best linear unbiased prediction (BLUP) of the random slope deviation. Participants contributed to the model when at least one non-missing episodic memory measurement was available; no minimum number of post-baseline assessments was required. Accordingly, slope estimates for participants without a post-baseline episodic memory assessment were predominantly informed by the population model and the estimated random-effects structure.

### Serological assessment of antiviral antibodies

Antibodies against widespread herpesviruses were measured in serum by enzyme-linked immunosorbent assay (ELISA) or indirect immunofluorescence at the Institute of Medical Virology, University of Zurich. Quantitative measurements were available for immunoglobulin G (IgG) antibodies against cytomegalovirus (CMV), herpes simplex virus (HSV) types 1 and 2, and varicella zoster virus (VZV). CMV IgG was reported in AE/ml with a positivity threshold of ≥6; HSV-1 and HSV-2 IgG were reported as signal-to-cut-off ratios (S/CO), with a positivity threshold of >1; and VZV IgG was reported in mIU/ml with a positivity threshold of >100. Quantitative CMV IgG values in seronegative participants were reported as ≤4 AE/ml. Antibody titres were log-transformed when included as continuous covariates in statistical models.

### Plasma AD biomarkers

Blood of study participants was collected in ethylenediaminetetraacetic acid (EDTA) tubes (Vacutainer, BD), inverted ten times and centrifuged at 1,620 × g for 12 min at 6 °C. Plasma supernatant was aliquoted and stored at −80°C until analysis. For study population 1, biomarker measurements were performed in the laboratories of H. Zetterberg and K. Blennow at Sahlgrenska University Hospital, Mölndal, Sweden. Plasma Aβ42, Aβ40, neurofilament light chain (NfL) and glial fibrillary acidic protein (GFAP) were quantified using a customized Neurology 4-Plex A Simoa assay (Quanterix). Plasma phosphorylated tau at threonine 217 (p-tau217) was measured using an in-house University of Gothenburg assay.^58^ Fluid biomarker levels below the limit of quantification (LOQ) were set at LOQ/2 according to standard approaches for left-censored LOQ data.^59^ For study population 2, plasma Aβ42, Aβ40 and total tau were measured by Quanterix on a Simoa HD-X instrument using the Human Neurology 3-Plex A assay. Plasma phosphorylated tau at threonine 181 (p-tau181) was measured using a Simoa p-tau181 immunoassay.

### C-reactive protein (CRP) assessment

Serum CRP concentrations were measured as part of routine clinical diagnostics in externally certified laboratories (Unilabs, Switzerland) to identify acute infection or inflammation.

### In vitro antigen stimulation assays

Cryopreserved PBMCs from study population 2 were thawed at 37 °C and washed twice in complete RPMI 1640 medium (Sigma-Aldrich) supplemented with 2 mM glutamine, 1% (v/v) non-essential amino acids, 1% (v/v) sodium pyruvate, penicillin (50 U/ml), streptomycin (50 μg/ml; all from Invitrogen) and 5% (v/v) heat-inactivated human serum (Swiss Red Cross). Monocytes were enriched by positive selection using CD14-coated microbeads (Miltenyi Biotec). From the CD14-depleted fraction, live singlets were selected using LIVE/DEAD Aqua viability dye (Invitrogen), followed by gating on CD3+CD19− T-cells. Memory CD4+ and CD8+ T-cells were then isolated by fluorescence-activated cell sorting (FACS) to >98% purity on a FACSAria Fusion (BD Biosciences) by excluding CCR7+CD45RA+ naïve-phenotype T-cells. In addition, CD25-bright, CD14+ and CD56+ cells were excluded. CD8+ cells were excluded during CD4 T-cell isolation, whereas CD4+ cells were excluded during CD8 T-cell isolation. All fluorochrome-conjugated mouse monoclonal antibodies used for sorting are reported in **Supplementary Table 3**. Cells were stained on ice for 15-20 min before sorting.

Following sorting, memory CD4 and CD8 T-cells were labelled with 5 μM carboxyfluorescein succinimidyl ester (CFSE, CellTrace, Invitrogen, ThermoFisher) and co-cultured with autologous monocytes irradiated with a total dose of 45 Gy at a T-cell-to-monocyte ratio of 2:1. Before co-culture, monocytes were either left untreated or pulsed for 1 h with selected peptide pools spanning the Aβ N-terminal, mid-sequence or C-terminal regions at 3 μg/ml per peptide (**Supplementary Table 2**, custom peptide synthesis service, Mimotopes/Biomatik). Inflexal V (5 μg/ml, Abbott) or an Epstein-Barr virus (EBV) human leukocyte antigen (HLA) class I peptide pool comprising 46 peptides (1 μg/ml, PEPotec, ThermoFisher) was used as a control antigen. Because of the limited numbers of memory CD8 T-cells available, CD8 T-cell responses were assessed only against a 1:1:1 mixture of the three Aβ peptide pools and the corresponding control conditions.

### Flow-cytometric assessment of antigen-specific T-cell responses

After six days of culture, cells were stained with antibodies against activation markers CD25 and ICOS (details in **Supplementary Table 3**). Cells were acquired on a LSR Fortessa flow cytometer (BD Biosciences) using BD FACSDiva software version 9.0, flow cytometry data were analysed using FlowJo version 10.8.1. After debris and doublet exclusion, separately cultured CD4 and CD8 antigen-responsive T-cell fractions were identified by CFSE dilution and activation-marker upregulation. Proliferating cells were defined as CFSE-low events relative to the undivided CFSE-high population present in unstimulated control cultures. CD25 and ICOS expression was assessed within the CFSE-low compartment. Gates were established using both unstimulated and positive controls, and applied consistently across samples.

A T-cell response was classified as positive only when both of the following criteria were met: (I) a stimulation index of ≥2, calculated as the percentage of CFSE-low cells in cultures containing antigen-pulsed autologous monocytes divided by the percentage of CFSE-low cells in matched cultures containing unpulsed autologous monocytes; and (II) a Δ value of ≥1.5%, calculated as the difference between the aforementioned two conditions. These thresholds were selected based on previous observations across negative and positive samples assessed using *ex vivo* T-cell stimulation assays with self-antigens.^60^

### Sample preparation and staining for mass cytometry (CyTOF)

Cryopreserved PBMCs were thawed at 37 °C and washed in RPMI 1640 medium (Sigma) containing 10% (v/v) heat-inactivated fetal bovine serum (FBS, Gibco) and 1% (v/v) GlutaMAX (Gibco). Approximately 1 × 10^6^ PBMCs per participant were transferred to a 96-well plate, washed in phosphate-buffered saline (PBS) and incubated with Cell-ID ^198^Pt-cisplatin (Fluidigm, diluted 1:10,000 in PBS) for 10 min at room temperature (RT) to identify dead cells. Cells were incubated with Human TruStain FcX Fc-receptor-blocking reagent (BioLegend) diluted 1:20 in Cell Staining Buffer (CSB, Fluidigm) for 5 min at RT. Customized live-cell barcoding was performed for 20 min using combinations of anti-human CD45 antibodies conjugated to different metal tags in a 10-choose-4 combinatorial design, allowing up to 210 unique barcode combinations. After barcoding, samples were washed, pooled and incubated for 20 min at RT with a mass-cytometry antibody panel containing major PBMC lineage markers and T-cell state and activation markers. Barcode and surface-staining antibodies are listed in **Supplementary Table 1**. After washing in CSB, cells were fixed with 1.6% (v/v) formaldehyde in CSB and stored overnight at 4°C. Stained cells were aliquoted in batches of approximately 20 million cells and frozen in 10% (v/v) dimethyl sulfoxide (DMSO) in FBS at −80 °C. For acquisition, frozen aliquots were thawed in pre-warmed PBS and incubated with Cell-ID Intercalator-Ir, diluted 1:5,000 in Maxpar Fix and Perm Buffer (Fluidigm), for 1 h at RT. Cells were washed in deionized metal-free water and Cell Acquisition Solution (CAS, Fluidigm) and adjusted to 1.2 × 10^6^ cells/ml in CAS containing EQ Four Element Calibration Beads (Fluidigm) at a 1:10 (v/v) dilution. Samples were acquired using a Helios CyTOF2 mass cytometer (DVS Sciences, Fluidigm).

### Mass cytometry data pre-processing

Mass-cytometry data were processed in R using *Premessa*, *CATALYST* and associated Bioconductor packages.^61–63^ Acquisition fragments were inspected using the DNA-intercalator signal against acquisition time, and intervals showing unstable acquisition were excluded. Signal intensities were normalized using EQ Four Element Calibration Beads (Fluidigm). Spillover affecting highly expressed metal-labelled barcode channels was estimated using single-stain controls and corrected by non-negative-least-squares compensation.^64^ Pooled events were computationally debarcoded and assigned to individual participants, and acquisition fragments corresponding to the same participant were concatenated. The CU, MCI and OO samples were acquired in one principal experiment. The Y samples were acquired in a subsequent experiment, processed using the same general workflow and harmonized with the principal dataset. Marker expression distributions were aligned and batch effects were corrected using *cyCombine*.^47^

### Mass cytometry data analysis

Mass-cytometry analyses were performed using *CATALYST*, *FlowSOM*, *ConsensusClusterPlus* and *diffcyt* packages.^62,65^ Lineage markers were used for unsupervised FlowSOM clustering with an 8 × 8 grid. Resulting nodes were manually annotated or merged into 24 biologically defined landmark populations based on canonical lineage and differentiation-marker expression.^32,66^ Landmark frequencies were calculated as percentages of total PBMCs for each participant. Median marker expression within landmark populations was used for heatmap visualization. Uniform manifold approximation and projection (UMAP) was calculated using a subsample of 10,000 cells, up to 5 million cells were displayed across study participants. High-resolution FlowSOM nodes were projected onto scaffold maps using the manually annotated landmark populations as reference nodes. Force-directed graphs were generated using the *vite* package and visualized using Fruchterman-Reingold layouts implemented in *igraph* and *ggraph*.^67,68^ The same cluster definitions were used across diagnostic and Aβ groups. Differential-state (DS) analyses were performed on cell-state markers using the diffcyt-DS-limma method.^65^ A differential-state feature was defined as a cluster-marker combination showing altered median marker expression between the compared participant groups. Models included sex and CMV IgG as covariates. Raw percentage changes in marker expression were used for graphical presentation, whereas statistical significance was determined from Benjamini-Hochberg false-discovery-rate-adjusted (BH FDR) *P* values.

### Statistical analysis

Statistical analyses were performed in R using *rstatix* and *stats* packages.^69^ All tests were two-sided. Available complete observations were used for each analysis. Exact sample sizes are reported in the figures, tables and source data.

#### Characterization of study participant subgroups

Continuous cohort characteristics were summarized as mean ± standard deviation (SD) or median and interquartile range (IQR), depending on skewed variables. Normally distributed variables across multiple groups were compared using one-way ANOVA followed by pairwise Welch tests, whereas non-normally distributed variables across multiple groups were compared using Kruskal-Wallis tests followed by Dunn’s tests. Categorical variables were compared using χ² or Fisher’s exact tests. Pairwise comparisons in Extended Data Table 1 and 2 were performed against the Aβ-negative CU reference group, with BH FDR correction within each table row.

#### Baseline biomarker correlations

Pairwise associations among baseline AD biomarkers, imaging measures and longitudinal outcomes were assessed using Spearman’s rank correlations. *P* values were adjusted across all tested correlations within each diagnostic group using the BH FDR procedure.

#### Association of landmark cluster frequencies with serology and sex

Associations of viral serology and sex with landmark-cell frequencies were estimated using linear regression. Viral-antibody effects were adjusted for age and sex, whereas sex effects were adjusted for age. Quantitative viral IgG titres were log-transformed. Regression coefficients for viral titres represent the change in landmark frequency, expressed in percentage points, per unit increase in log-transformed titre. Regression coefficients for sex represent the difference in landmark frequency between male and female participants.

#### Healthy-aging immune-cell analyses

Healthy-aging analyses included Y, Aβ-negative CU and Aβ-negative OO participants. Raw immune-cell frequencies and marker-expression values are displayed in violin plots. Because clinical covariates were unavailable for the Y group, a hybrid adjustment approach was used: values from CU and OO participants were residualized using linear models including sex and CMV IgG titre, whereas Y values remained unadjusted. Pairwise group differences were assessed using Dunn’s tests. *P* values were adjusted across pairwise comparisons within each landmark population using the BH FDR procedure.

#### Immune cell state analyses with *diffcyt*

Differential-state (DS) analyses evaluated disease-relevant contrasts and included sex and CMV IgG as covariates. *P* values were adjusted within each contrast using the BH FDR procedure, with FDR-adjusted P < 0.05 considered significant. Raw percentage changes were used to show the magnitude and direction of DS effects. For ranked displays, the feature with the greatest absolute raw percentage change was retained when duplicate features occurred, and features were ordered according to their absolute percentage change.

#### Linear regression screening and recursive feature elimination

Associations between individual immune features and continuous cerebral Aβ load were screened using linear regression. Aβ PET SUVR parameters were natural-log transformed, immune markers were standardized to one SD, and models were adjusted for sex and CMV IgG titre. *P* values were adjusted across the tested immune features using the BH FDR procedure. To determine whether immune markers predicted continuous cerebral Aβ load, random-forest recursive feature elimination (RF-RFE) was performed using fixed tenfold cross-validation (CV) folds. Feature selection and model fitting were conducted within the CV framework. Predictive performance was evaluated using pooled out-of-fold predictions and quantified by the root-mean-square error (RMSE) and the squared Pearson correlation between observed and predicted natural-log-transformed Aβ PET SUVR. For the RF-RFE-selected marker subset, permutation importance was quantified as the percentage increase in out-of-bag mean squared error following permutation of each selected feature, with values summarized across cross-validation resamples.

#### Logistic-regression models

Logistic-regression models were used to classify cerebral Aβ status. Models were fitted separately within the CU, MCI and OO groups using participants with complete data for all predictors included in the models being compared. A reference model included baseline age, sex and *APOE* ε4 carrier status. The plasma AD biomarker model additionally included the plasma Aβ42/Aβ40 ratio and p-tau217. The immune model additionally included inducible T-cell co-stimulator (ICOS) expression in CD4 effector-memory (CD4 EM), CD8 effector-memory (CD8 EM) and CD8 central-memory (CD8 CM) T-cell populations. Receiver operating characteristic (ROC) curves and areas under the curve (AUCs), with 95% confidence intervals (CIs), were calculated from model-predicted probabilities. The immune and plasma AD biomarker models were compared using paired DeLong tests.

#### Longitudinal episodic-memory models

Longitudinal episodic-memory trajectories were analysed separately in the CU, MCI and OO groups using LMMs fitted with the *nlme* package. Models included follow-up time in years from baseline, baseline Aβ PET SUVR and their interaction. Baseline age, sex and years of education were included as covariates together with their respective interactions with time. Participant-specific random intercepts and random slopes for time were included. Term of interest was the time × Aβ PET SUVR interaction, representing the association between baseline cerebral Aβ burden and subsequent episodic memory change. *P* values for this interaction were adjusted globally across the three diagnostic groups using BH FDR procedure. Model-adjusted marginal episodic memory trajectories and pointwise 95% CIs were generated from the fixed-effects component of each group-specific LMM using the *effects* package. Years from baseline and Aβ PET SUVR were each evaluated at five automatically selected, approximately equally spaced values spanning their ranges in the corresponding group-specific model data, with values rounded by the package for graphical display. The resulting fitted values were connected to visualize the estimated time × Aβ PET SUVR interaction. A common time axis spanning the overall observed follow-up range was used across the CU, MCI and OO panels.

#### Immune-marker interaction models for longitudinal outcomes

Linear regression models were used to determine whether baseline immune-marker expression modified associations between cerebral Aβ load and hippocampal volume change (atrophy), WMH volume change or episodic memory change over time. Analyses were performed separately within the CU, MCI and OO groups. Immune markers were standardized to a mean of zero and standard deviation of one. For each outcome, three models were examined: a base model (‘Model 1’) containing covariates and Aβ PET SUVR; an additive model (‘Model 2’) additionally containing the standardized immune marker; and an interaction model (‘Model 3’) containing the Aβ PET SUVR × immune-marker interaction. Models on hippocampal volume change were adjusted for baseline age, sex and TIV. Models on WMH volume change were adjusted for baseline age and sex. Models on episodic memory change were adjusted for baseline age, sex and years of education. The interaction coefficient β represents the change in the association between Aβ PET SUVR and the longitudinal outcome per one-standard-deviation higher immune-marker expression.

*P* values for tested interaction terms were adjusted using BH FDR procedure within each diagnostic group. Significant interactions were visualized using model-based predictions at low (−1 standard deviation), mean and high (+1 standard deviation) immune-marker expression.

#### Longitudinal diagnostic-converter analyses

Baseline immune cluster frequencies and marker expressions were compared between longitudinally stable participants and participants showing persistent conversion from one diagnosis to another. For statistical testing, each immune outcome was residualized using a linear model including sex and qualitative CMV IgG status. The resulting residuals were compared using a two-sided Wilcoxon rank-sum test. Cliff’s delta was calculated as a non-parametric estimate of effect size; negative values indicate lower values in converters and positive values indicate higher values in converters. Reported *P* values therefore refer to tests performed on covariate-adjusted residuals.

### Data availability

The code generated during this study for data analysis is publicly available at Zenodo (CERN, European Organization for Nuclear Research) under https://doi.org/10.5281/zenodo.21777909.

### Ethics declaration

The participants included in the analyses participated in in-house cohort studies conducted by the ‘Center for Prevention and Dementia Therapy’ at the Institute for Regenerative Medicine at the University of Zurich, Switzerland. All study participants gave written informed consent. The cohort studies including the here shown assessments were approved by the local ethics committee (Kantonale Ethikkommission, Zurich, Switzerland) and conducted in accordance with their guidelines and the Declaration of Helsinki.^70^

## Results

### A deeply phenotyped study population capturing ageing and early AD pathology

We first characterized study population 1, an in-house monocentric study population, which comprised 200 participants aged 25 to 96 years at baseline. Participants were assigned to four groups: young adults (Y; n = 10), cognitively unimpaired older adults (CU; n = 95), participants with mild cognitive impairment (MCI; n = 47) and oldest-old participants aged ≥85 years without dementia (OO; n = 48) (**Fig. 1a**, **Table 1 and Extended Data Table 1**). The Y group did not undergo clinical examination or biomarker analysis but was used exclusively as young reference group to compare the immune architecture of older and younger subjects. In contrast, CU, MCI and OO participants underwent extensive clinical, neuropsychological, imaging and biomarker characterization at baseline, with longitudinal follow-up assessments for cognitive and MRI-based outcomes (**Fig. 1b**). Baseline cerebral Aβ load was assessed by [18F]-flutemetamol PET, and CU, MCI and OO participants were stratified into Aβ-negative (CU−, MCI− and OO−) or Aβ-positive (CU+, MCI+ and OO+) subgroups (**Extended Data Fig. 1a,b**). Additional baseline assessments included serological measurement of antibodies against widespread herpesviruses and plasma AD biomarker profiling.

**Fig. 1:**
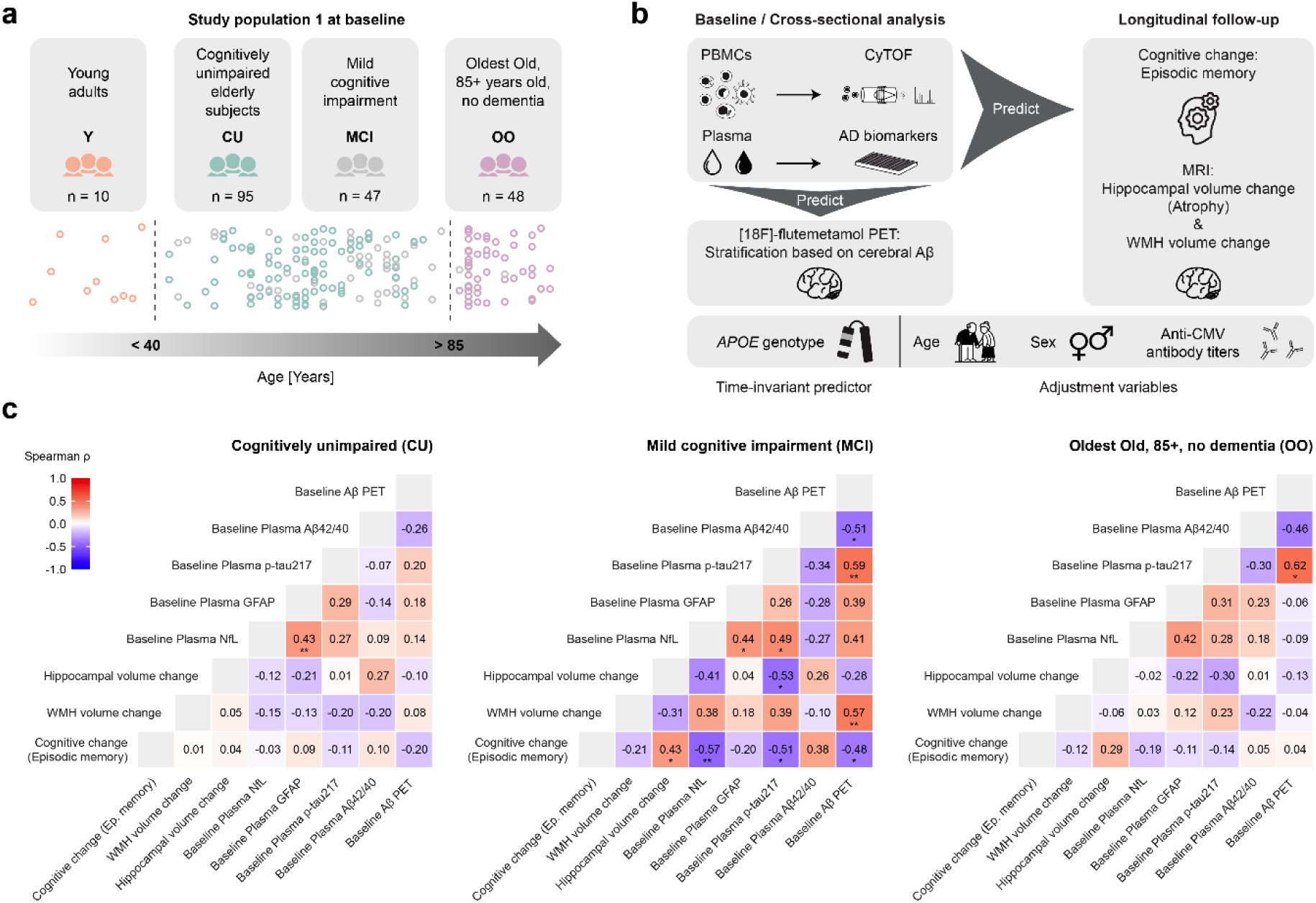
Study design and biomarker characterization of participant subgroups. **a**. Overview of the study population 1 at baseline. A total of 200 participants from a heterogeneous cohort spanning healthy aging and early Alzheimer’s disease (AD) were assigned to four subgroups based on age and cognitive status: young adults (Y; n = 10), elderly cognitively unimpaired participants (CU; n = 95), participants with mild cognitive impairment (MCI; n = 47) and a group of exceptional agers ≥85 years without dementia called ‘oldest-old’ (OO; n = 48). **b**. Schematic overview of baseline and longitudinal analyses. At baseline, peripheral blood mononuclear cells (PBMCs) were analyzed by CyTOF for immunological profiling. In the CU, MCI and OO groups, AD-related biomarkers were assessed using plasma AD biomarker measurements and [18F]-flutemetamol PET imaging for cerebral Aβ stratification. Longitudinal follow-up for up to 8 years included neuropsychological assessment of episodic memory, as well as MRI-based quantification of hippocampal volume change and white matter hyperintensity (WMH) volume change. Additional variables used in modelling included *APOE* genotype, age, sex and serum antibody titres against latent *Herpesviridae* such as Cytomegalovirus (CMV). **c**. Heatmaps show pairwise correlations among baseline AD biomarkers and longitudinal outcome measures in CU, MCI and OO participants. Panels display age- and sex-adjusted Spearman correlation coefficients (ρ). Colors represent the magnitude and direction of the correlation coefficient. Significance symbols indicate Benjamini-Hochberg false discovery rate (FDR)-adjusted *P* values, corrected across all pairwise correlations within each diagnostic group. * *P* < 0.05, ** *P* < 0.01.

Aβ42 was assessed as a standard plasma biomarker of cerebral Aβ pathology and normalized to Aβ40 to account for inter-individual and age-related differences in overall Aβ production and clearance. The plasma Aβ42/40 ratio inversely tracked cerebral Aβ status, with the strongest group-specific association observed in MCI participants (**Fig. 1c, Extended Data Fig. 1c,d**). Plasma p-tau217 was higher in Aβ-positive groups with substantial cerebral Aβ burden, particularly in MCI+ and OO+ (**Fig. 1c, Extended Data Fig. 1e**). Plasma levels of NfL and GFAP, reflecting neuroaxonal injury and astroglial activation in neurodegenerative disease,^6^ were elevated in MCI+ participants and in the OO groups, where higher age is likely to contribute to these increases (**Extended Data Fig. 1f,g**). NfL and GFAP were positively correlated with each other in CU and MCI participants (**Fig. 1c**).

MRI-based outcome measures were available at baseline and during longitudinal follow-up. We assessed hippocampal volume as a structural marker of neurodegeneration and cognitive decline, dependent or independent of AD pathology.^71,72^ Baseline hippocampal volume was lower and longitudinal volume change was more negative in the AD pathology-burdened MCI+ participants as well as in the OO− and OO+ groups, indicating increased hippocampal atrophy (**Extended Data Fig. 1h,i**). In MCI participants, hippocampal volume loss correlated with higher baseline plasma p-tau217 levels (**Fig. 1c**). In addition, we quantified white matter hyperintensities (WMHs), which reflect small-vessel-related brain injury and are associated with vascular risk factors relevant to AD.^73^ Baseline WMH volume was higher and longitudinal WMH volume change was more positive in MCI+ participants and in the OO− and OO+ groups, indicating increased WMH accumulation (**Extended Data Fig. 1j,k**). In the MCI group, longitudinal WMH volume change showed a positive correlation with baseline cerebral Aβ load (**Fig. 1c**).

Cognitive performance was assessed at baseline and at approximately annual neuropsychological follow-up visits. Global cognitive performance was summarized using the Mini-Mental State Examination (MMSE), which captured the expected reduction in global cognitive performance in MCI participants (**Table 1, Extended Data Table 1**). Domain-specific neuropsychological testing included language function, processing speed, executive function, episodic memory and visuospatial abilities. In the present study, we focused on an episodic memory composite score because episodic memory is among the earliest and most prominently affected cognitive domains in AD^45^ and is closely linked to hippocampal atrophy over time.^74^ Baseline episodic memory performance was lower and longitudinal episodic memory change was more negative in MCI participants and in the OO groups (**Extended Data Fig. 1l,m**). Particularly in the MCI group, episodic memory decline correlated with hippocampal atrophy and higher baseline levels of cerebral Aβ, plasma p-tau217 and plasma NfL (**Fig. 1c**).

The frequency of *APOE* ε4 carriers, the strongest common genetic risk factor for sporadic AD, was increased in the CU+ and MCI+ groups, but not in OO+ (**Table 1, Extended Data Fig. 1a**). Consistent with the association between *APOE* ε4 and cerebral Aβ pathology, *APOE* ε4 carriers showed higher cerebral Aβ load and lower plasma Aβ42/40 ratio than non-carriers in the pooled sample (**Extended Data Fig. 1n,o**).

Together, these data show that study population 1 spans a deeply phenotyped continuum from healthy young adulthood to cognitively unimpaired older age with or without preclinical AD pathology, early cognitive impairment and exceptional old age without dementia.

### High-dimensional mass cytometry defines immune-cell architecture across aging and study subgroups

To map the peripheral blood immune-cell architecture in study population 1, we performed high-dimensional mass cytometry (CyTOF) on baseline PBMC samples using a T-cell-biology-focused antibody panel comprising lineage, differentiation, activation and cell trafficking markers. Automated, unsupervised FlowSOM clustering based on lineage markers was used to define high-resolution immune-cell clusters, which were subsequently mapped onto manually annotated landmark clusters. Landmark cluster identities were assigned based on median marker-expression patterns and the annotated landmark clusters were organized into major immune cluster families, including B-cells, CD4 T-cells, CD8 T-cells, myeloid cells, innate lymphocytes, unconventional T-cells and plasmacytoid dendritic cells (**Fig. 2a**). The resulting immune-cell architecture was visualized as a scaffold map of total CD45+ cells, in which landmark clusters served as reference nodes to map unsupervised high-resolution FlowSOM clusters according to marker-expression similarity (**Fig. 2b**). A complementary UMAP representation of total PBMCs confirmed the separation of the major landmark cluster families across the high-dimensional marker space (**Extended Data Fig. 2a**). At the participant level, stacked family-composition plots showed inter-individual variability within diagnostic and Aβ subgroups, while also indicating broad age-associated shifts in immune-cell composition, particularly in the Y group (**Extended Data Fig. 2b**). This reference architecture provided the basis for subsequent analyses of landmark cluster frequencies and cell-state alterations across aging, Aβ pathology and cognitive status.

**Fig. 2:**
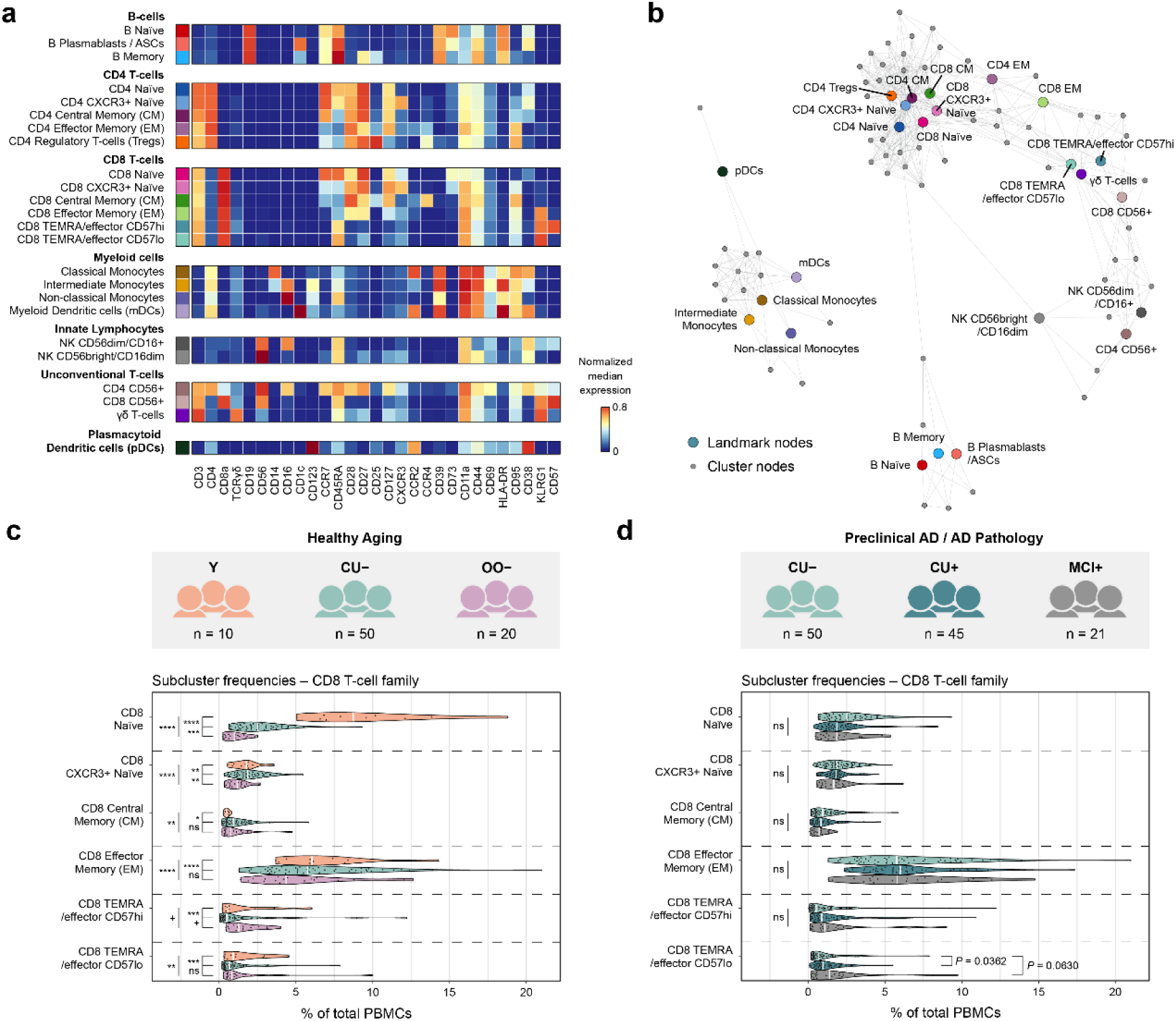
High-dimensional single-cell immune profiling identifies CD8 T-cell alterations during healthy aging and AD-associated pathology. **a**. Heatmap showing median marker expression across manually annotated immune landmark clusters. **b**. Scaffold map of total CD45+ cells. Each node represents a cell cluster, including landmark clusters (manually annotated, shown in colours corresponding to heatmap) and cluster nodes (unsupervised FlowSOM clusters, in grey). **c**. Healthy aging analysis including young adults (Y), PET Aβ-negative CU (CU−) and PET Aβ-negative OO (OO−). Participants were classified as cerebral Aβ-positive based on [18F]-flutemetamol PET Centiloid values > 12. Sideways violin plots showing frequencies of CD8 T-cell family landmark nodes as percentage of total PBMCs in the healthy aging analysis. White lines indicate medians. Pairwise group comparisons were performed using Dunn’s test on hybrid values; CU− and OO− values were residualized for sex and CMV antibody titre, whereas Y values remained raw. *P* values were adjusted using BH FDR across all pairwise comparisons within each landmark node. FDR-adjusted *P* values ≤ 0.1 are displayed. ^+^ *P* < 0.1, * *P* < 0.05, ** *P* < 0.01, *** *P* < 0.001, **** *P* < 0.0001. **d**. Preclinical AD / AD pathology analysis including CU−, PET Aβ-positive CU (CU+) and PET Aβ-positive MCI (MCI+). Sideways violin plots showing frequencies of CD8 T-cell family landmark nodes as percentage of total PBMCs in the Preclinical AD / AD pathology analysis. Pairwise comparisons were performed using Dunn’s test, with all groups adjusted for sex and CMV antibody titre. Testing was restricted to comparisons against the CU− reference group. *P* values were adjusted using BH FDR across the comparisons within each landmark node. FDR-adjusted *P* values ≤ 0.1 are displayed. Abbreviations: CU = cognitively unimpaired older adults; MCI = mild cognitive impairment; OO = oldest-old participants without dementia; PBMCs = peripheral blood mononuclear cells; BH FDR = Benjamini-Hochberg false discovery rate approach.

### Sex and herpesvirus serology are linked to peripheral immune-cell frequency shifts

Because age-associated immune remodelling can be influenced by sex and chronic viral exposure, we next assessed demographic and infection-related covariates of landmark cluster frequencies in CU, MCI and OO participants. Serum IgG titres against common, often latent herpesviruses, including cytomegalovirus (CMV), herpes simplex virus 1 (HSV-1), HSV-2 and varicella zoster virus (VZV), were analysed together with sex in landmark-cluster-wise linear models (**Extended Data Fig. 2c**).

CMV IgG titres showed the strongest associations with peripheral immune-cell composition. After adjustment for age and sex, higher CMV IgG titres were associated with increased frequencies of differentiated T-cell populations, most notably CD4 effector memory (EM) T-cells, CD8 EM T-cells, CD8 TEMRA/effector subsets, CD8 CD56+ T-cells and γδ T-cells, and with lower frequencies of CD4 central memory (CM) T-cells (**Extended Data Fig. 2c**). In contrast, HSV-1, HSV-2 and VZV IgG titres showed more limited associations with landmark cluster frequencies. Sex was also associated with immune-cell composition, with male participants showing higher classical monocyte frequencies and lower CD4 T-cell frequencies than female participants, consistent with known sex-related differences in peripheral immunity.^75^ Together with previously reported sex differences and herpesvirus associations in AD pathogenesis,^17,76–79^ these observed associations with peripheral immune-cell frequencies supported adjustment for sex and CMV IgG titre in subsequent analyses.

### Healthy aging and Aβ pathology converge on CD8 T-cell compartment changes

To identify age-associated immune cell differences independently of detectable cerebral Aβ pathology, we first compared Y participants with Aβ-negative CU and Aβ-negative OO participants across a healthy aging continuum. Within the CD8 T-cell family, healthy aging was associated with a redistribution of landmark cluster frequencies, most prominently including reductions in CD8 naïve and CD8 EM landmark clusters (**Fig. 2c**). Age-associated remodelling was not restricted to the CD8 T-cell compartment. Extended family-level analyses showed additional changes across CD4 T-cell, B-cell, myeloid-cell, innate-lymphocyte, unconventional T-cell and plasmacytoid dendritic cell (pDC) families, including reduced CD4 naïve T-cells and increased CD4 regulatory T-cells (Tregs) and classical monocytes (**Extended Data Fig. 2d**). These age-associated differences were observed after adjustment for sex and CMV IgG titre in participants with available covariate data.

We next asked whether early AD-pathology was associated with CD8 T-cell compartment changes. In a comparison of CU−, CU+ and MCI+ participants, representing potential preclinical AD and early AD pathology, CD8 T-cell landmark frequencies showed evidence of remodelling, with increased frequency of antigen-experienced, differentiated CD8 TEMRA/effector cells in the Aβ-positive CU+ and MCI+ groups compared with CU− participants (**Fig. 2d**). These results were consistent with earlier findings of changes within the CD8 T-cell compartment.^32,33^ Outside the CD8 T-cell family, frequency changes in this early AD-pathology comparison were observed in the CD4 T-cell family, with increased frequency of CXCR3+ naïve CD4 T-cells; no other landmark family showed significant differences in this setting (**Extended Data Fig. 2e**).

Together, these frequency-level analyses identified CD8 T-cell remodelling as a shared feature of healthy aging and AD-pathology-associated disease stages.

### Aβ pathology is associated with ICOS-centred T-cell state changes

Having identified CD8 T-cell compartment remodelling at the frequency level, we next investigated whether Aβ pathology was associated with altered immune-cell states within landmark clusters. To this end, we used the diffcyt framework to perform differential-state analyses of surface marker expression within immune-cell landmark clusters.^65^ While lineage markers were used to define the immune-cell clustering structure, activation, differentiation, trafficking, co-signalling and regulatory markers were tested as cell-state features. We focused on three prespecified AD-relevant contrasts: CU+ versus CU−, representing preclinical AD; MCI+ versus CU−, representing AD pathology progression; and MCI+ versus MCI−, representing AD versus non-AD. All differential-state models were adjusted for sex and CMV IgG titre. Across these contrasts, Aβ pathology was associated with recurrent T-cell state changes centred on the surface expression of the inducible co-stimulatory marker ICOS. In the preclinical AD contrast, CU+ participants showed increased ICOS expression across multiple CD4 and CD8 T-cell landmark clusters, including memory and CXCR3+ naïve clusters, compared with CU− participants (**Fig. 3a**). In the AD-pathology progression contrast, MCI+ participants showed a related pattern of increased ICOS expression on T-cell subsets compared with CU− participants, including CD4 and CD8 memory and TEMRA/effector compartments (**Fig. 3b**). In the AD versus non-AD contrast, comparing MCI+ with MCI− participants, differential-state changes again involved ICOS-expressing T-cell features, most prominently in memory, CD4 Treg and CD8 TEMRA/effector clusters (**Fig. 3c**). Additional disease-relevant differential-state hits were observed in B-cell and unconventional T-cell cluster families (**Extended Data Fig. 3a,b**). Next, we ranked the top ten diffcyt features by absolute raw percentage change while retaining only the contrast with the largest effect for each cluster-marker feature (**Fig. 3d**). This ranking highlighted ICOS as the dominant recurrent marker among diagnosis- and Aβ-associated immune-cell state changes, including ICOS expression in CD8 TEMRA/effector, CD4 and CD8 EM, CD4 and CD8 CM, CD4 Treg, and γδ T-cell compartments. These findings indicated that Aβ-associated immune-cell state alterations were not confined to a single T-cell subset, but converged on an ICOS-centred activation state across multiple T-cell compartments.

**Fig. 3:**
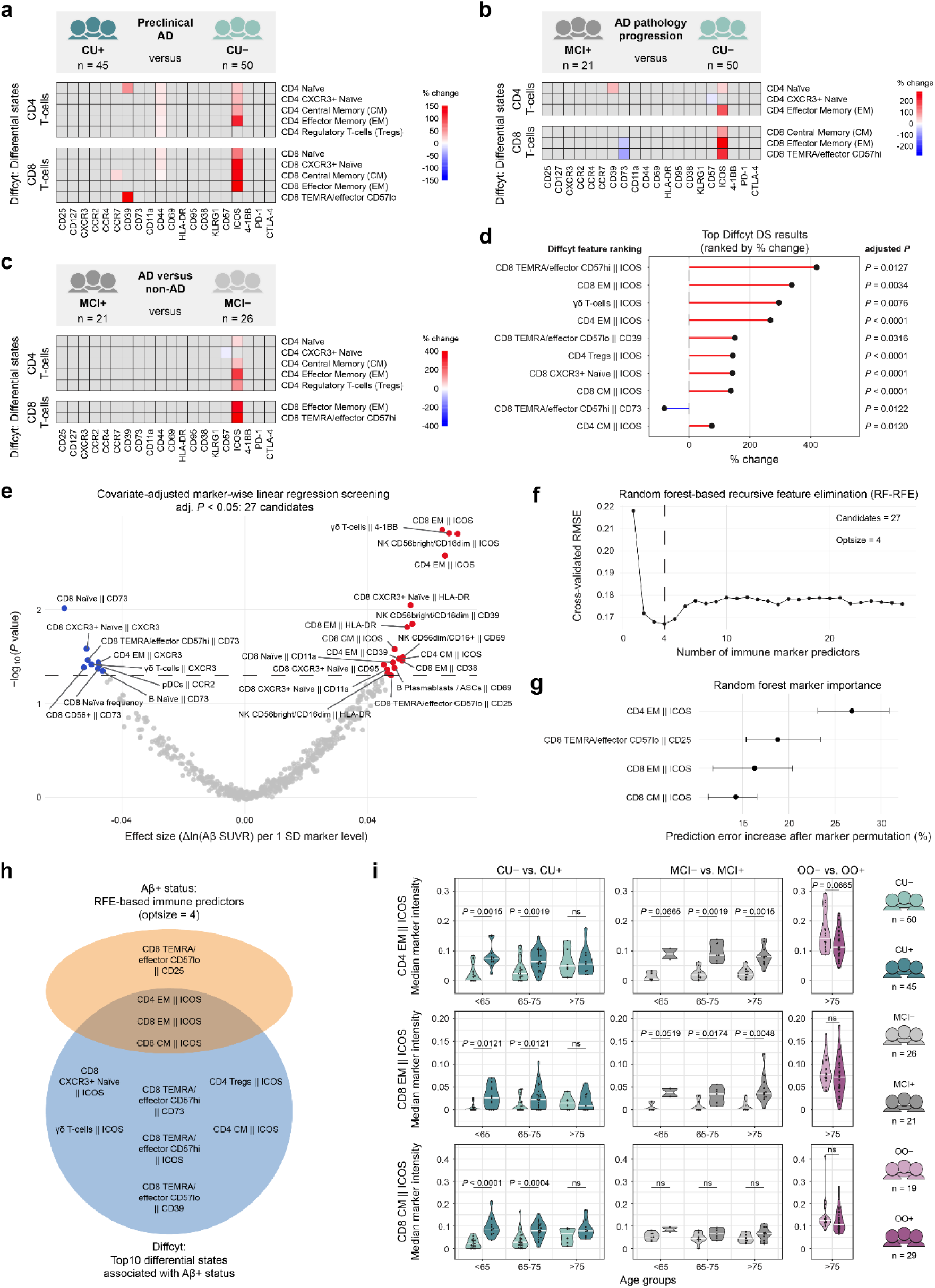
Immune cell state analysis identifies ICOS expression in T-cell subsets associated with Aβ pathology. **a-c**. Diffcyt differential state analyses showing AD-relevant immune marker changes within landmark clusters. Heatmaps show significant cluster-marker features for comparisons between PET Aβ-positive and Aβ-negative CU (CU+ versus CU− = preclinical AD; **a**), PET Aβ-positive MCI and PET Aβ-negative CU (MCI+ versus CU− = AD pathology progression; **b**), and PET Aβ-positive versus PET Aβ-negative MCI (MCI+ versus MCI− = AD versus non-AD; **c**). Analyses were adjusted for sex and CMV IgG. Colours indicate raw percentage change in marker expression. Grey tiles indicate features not passing the BH FDR-adjusted threshold of *P* < 0.05, and landmark clusters without significant hits are not shown. **d**. Ranking of the top disease-relevant Diffcyt differential state results across the preclinical AD, AD pathology and AD versus non-AD contrasts. For each cluster-marker feature, only the contrast with the largest absolute raw percentage change was retained to avoid duplicate representation across related contrasts. Features were ranked by absolute raw percentage change and the top ten are shown. Line colour indicates the direction of change, as in a–c. Labels indicate BH FDR-adjusted *P* values. **e**. Marker-wise linear regression screening for association with cerebral PET Aβ load. Each point in the volcano plot represents one immune marker tested in a linear model with cerebral Aβ load as outcome, and sex and CMV IgG titre as covariates. X-axis shows marker effect size, expressed as change in PET Aβ load, Δln(Aβ SUVR), per 1 SD higher immune marker level. Y-axis shows transformed BH FDR-adjusted *P* values. Dashed horizontal line indicates BH FDR-adjusted significance threshold of *P* < 0.05. Significant positive and negative associations are shown in red and blue, respectively (total = 27); non-significant markers are shown in grey. **f**. Random forest recursive feature elimination (RF-RFE) performance curve for prediction of cerebral Aβ load from the significant immune marker set identified in panel e. Curve shows 10-fold cross-validated root mean squared error (RMSE) as a function of the number of retained immune marker predictors. Vertical dashed line indicates the optimal subset size, defined as number of predictors yielding the lowest cross-validated RMSE. **g**. Random forest permutation importance for the RF-RFE-selected immune marker subset. Importance reflects the increase in mean squared prediction error after permutation of each marker. Higher values indicate a greater contribution to prediction. Points show mean importance values, and horizontal error bars indicate the 2.5th–97.5th percentile range across cross-validation resamples. **h**. Overlap between the top ten Aβ pathology-associated Diffcyt differential state features and the RF-RFE-selected immune predictors, shown as a Venn diagram. **i**. Aβ-negative versus Aβ-positive marker intensities for the three overlapping Aβ pathology-associated immune markers across diagnostic and age strata. Marker intensities are shown as median arcsinh-transformed CyTOF signal. CU and MCI participants were stratified into age groups based on the 25th and 75th percentiles of baseline age among the CU and MCI groups; OO participants are shown as a singular group. *P* values were calculated using Wilcoxon rank-sum tests within each diagnosis-by-age stratum and adjusted using BH FDR within each marker. Abbreviations: CU = cognitively unimpaired older adults; MCI = mild cognitive impairment; BH FDR = Benjamini-Hochberg false discovery rate approach; SD = standard deviation; RF-RFE = Random forest recursive feature elimination; RMSE = root mean squared error; OO = oldest-old participants without dementia.

We next tested whether the immune-cell state findings identified in categorical disease contrasts were also associated with continuous cerebral Aβ load. In marker-wise linear regression models with cerebral Aβ load as outcome and sex and CMV IgG titre as covariates, 27 immune-marker features were associated with cerebral Aβ load (**Fig. 3e**). Positively associated features again included several ICOS-expressing T-cell landmark clusters, whereas negatively associated features predominantly involved CD73 and CXCR3 markers across various T-cell compartments. To identify an optimal marker set predictive of continuous Aβ load, we then applied random-forest recursive feature elimination (RF-RFE) to the 27 Aβ-load-associated immune-marker candidates. RF-RFE selected a four-marker model with the lowest cross-validated prediction error (root mean squared error, RMSE) (**Fig. 3f**). Cross-validated model performance was evaluated using out-of-fold predictions, yielding RMSE = 0.167 and R² = 0.347, with R² calculated as the squared correlation between observed and predicted cerebral Aβ load (**Extended Data Fig. 3c**). Marker importance within the RF-RFE-selected model was assessed by permuting each immune-marker feature and quantifying the resulting increase in prediction error. This permutation-importance analysis ranked CD4 EM || ICOS as the strongest contributor, followed by CD8 TEMRA/effector CD57lo || CD25, CD8 EM || ICOS and CD8 CM || ICOS (**Fig. 3g**). Finally, we compared the top ten diffcyt differential-state features with the RF-RFE-selected predictors. Three ICOS-expressing memory T-cell clusters overlapped between both approaches: CD4 EM || ICOS, CD8 EM || ICOS and CD8 CM || ICOS and were selected for subsequent analyses (**Fig. 3h**). Stratified visualization across diagnostic and age groups showed higher ICOS marker intensities in cerebral Aβ-positive compared with Aβ-negative participants across most CU and MCI strata (**Fig. 3i**). This Aβ-related difference was attenuated in CU participants aged >75 years and was absent or reversed in OO participants, where already Aβ-negative individuals showed higher ICOS marker intensities. Together, these complementary analyses identified ICOS-centred T-cell state changes as a prominent immune correlate of cerebral Aβ pathology.

### ICOS-expressing memory T-cell subsets support Aβ classification and relate to hippocampal atrophy

We next asked whether ICOS surface expression on CD4 EM, CD8 EM and CD8 CM T-cell clusters supported classification of cerebral Aβ status beyond established demographic, genetic and plasma biomarker information. Within CU, MCI and OO participants, we compared logistic regression models predicting cerebral Aβ status using a reference model including age, sex and *APOE* ε4 carrier status; a plasma AD biomarker model that added plasma Aβ42/Aβ40 ratio and p-tau217 to the reference model; and an immune-marker model that added median ICOS signal intensities in CD4 EM, CD8 EM and CD8 CM T-cells to the reference model. In CU participants, the immune-marker model showed better discrimination of cerebral Aβ status than both the reference and plasma AD biomarker models, with AUCs of 0.728, 0.784 and 0.887 for the reference, plasma AD and immune-marker models, respectively (paired DeLong *P* = 0.0395 for immune-marker versus plasma AD biomarker model) (**Fig. 4a**). In MCI participants, the immune-marker model also showed higher Aβ classification performance, with AUCs of 0.736, 0.876 and 0.982 for the reference, plasma AD and immune-marker models, respectively, with weaker evidence of improvement over the plasma AD biomarker model (paired DeLong *P* = 0.0878) (**Fig. 4b**). In OO participants, the immune-marker model did not improve Aβ classification over the plasma AD biomarker model, with AUCs of 0.575, 0.825 and 0.707 for the reference, plasma AD and immune-marker models, respectively (paired DeLong P = 0.1858) (**Fig. 4c**). Thus, ICOS expression on memory T-cell subsets supported cerebral Aβ classification in CU and MCI participants, but not in the OO group.

**Fig. 4:**
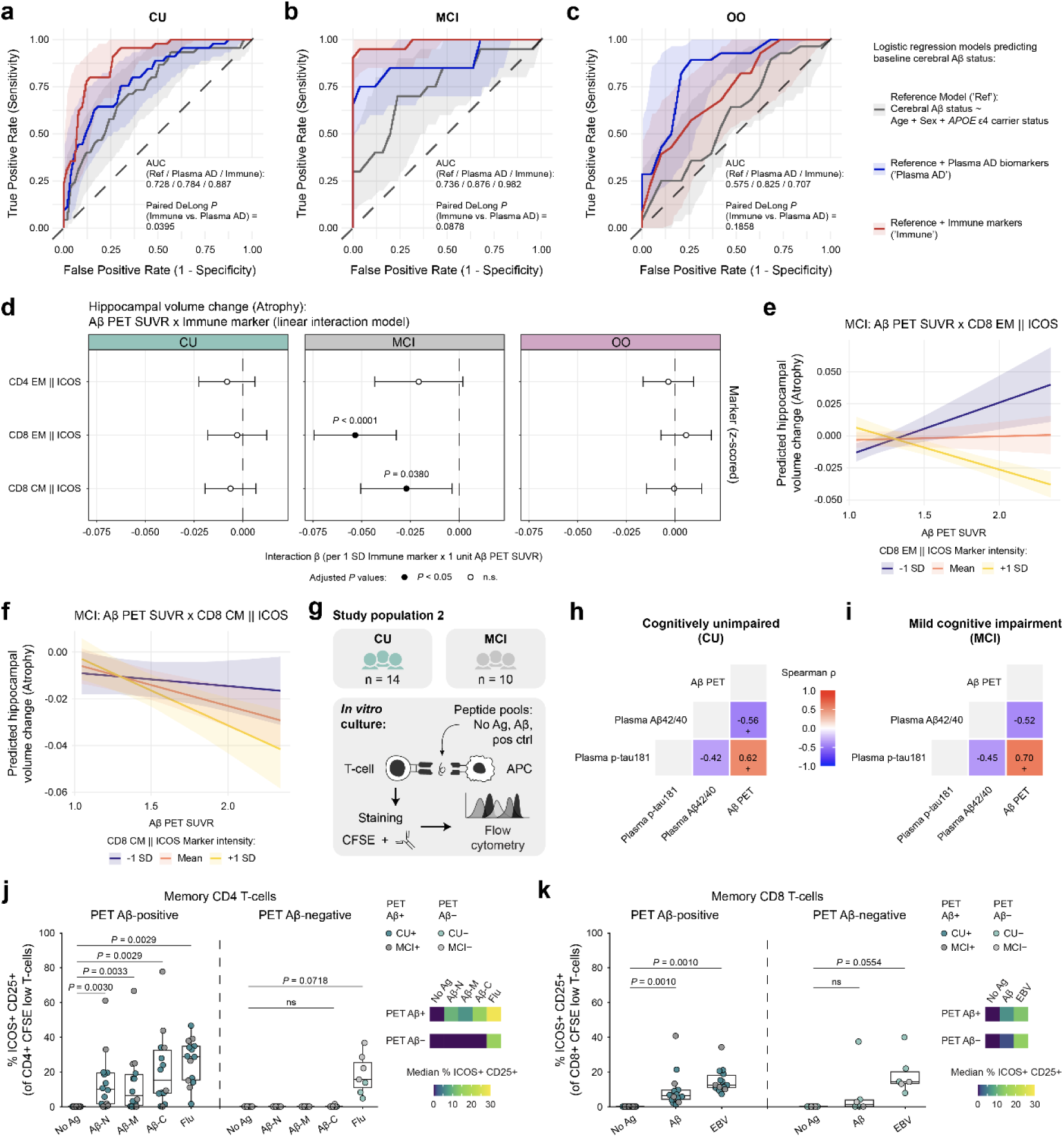
ICOS expression in memory T-cell subsets supports Aβ classification and aligns with Aβ-reactive T-cell responses. **a-c**. Receiver operating characteristic (ROC) curves from logistic regression models predicting cerebral Aβ status in CU (a), MCI (b) and OO (c). Three models were compared: a reference model including age, sex and *APOE* ε4 carrier status (= Ref); a reference + plasma AD biomarker model additionally including plasma Aβ42/Aβ40 ratio and plasma p-tau217 levels (= Plasma AD); and a reference + immune marker model additionally including median ICOS signal intensities in peripheral CD4 EM, CD8 EM and CD8 CM T-cells (= Immune). Shaded areas indicate 95% confidence intervals. Diagonal dashed line indicates random classification. AUC values and paired DeLong test *P* values comparing Ref + Immune model with Ref + Plasma AD model are shown within each panel. **d**. Linear interaction models testing whether immune-marker expression modified the association between cerebral Aβ load and longitudinal hippocampal volume change. Models were adjusted for baseline age, sex and TIV at baseline, and included Aβ PET SUVR, immune-marker expression and their interaction. Forest plots show Aβ SUVR × immune-marker interaction β estimate and 95% confidence interval; β reflects the change in Aβ PET-hippocampal volume change association per 1 SD higher immune-marker expression. Filled circles indicate BH FDR-adjusted *P* < 0.05 within each diagnostic group. **e,f**. Model-based prediction plots for significant interactions in MCI participants. Predicted hippocampal volume change is shown across the observed range of Aβ SUVR at low (−1 SD), mean and high (+1 SD) immune-marker expression, illustrating how immune-marker levels modify the association between cerebral Aβ load and hippocampal atrophy. Predictions represent conditional estimates from an unconstrained linear model. **g**. Study population 2: Independent confirmation study and *in vitro* antigen-stimulation workflow. Memory CD4+ and CD8+ T-cells from CU (n = 14) and MCI (n = 10) participants were sorted, CFSE-labelled and stimulated *in vitro* for 6 days with Aβ-derived peptide pools or control antigens, followed by antibody staining and flow cytometry analysis. **h,i**. Spearman correlation heatmaps showing associations among available AD biomarker measures in the independent confirmation cohort, shown for CU (h) and MCI (i). *P* values were adjusted using BH FDR within each diagnostic group. + P < 0.1. **j,k**. Frequencies of ICOS+CD25+ cells among CFSElow memory CD4 (j) and CD8 (k) T-cells after 6 days of *in vitro* culture under the indicated stimulation conditions. CD4 T-cell stimulation conditions included no antigen (No Ag), Aβ-N, Aβ-M, Aβ-C and influenza virus antigens (Flu) as positive control. Aβ-N, Aβ-M and Aβ-C correspond to peptide pools derived from the N-terminal, mid-sequence and C-terminal regions of Aβ1-42 peptide, respectively. CD8 T-cell stimulation conditions included No Ag, pooled Aβ-derived peptides and EBV antigens as positive control. Participants were grouped by cerebral Aβ PET status; point colours indicate the underlying clinical subgroup. Each point represents one participant; boxplots show median and interquartile range. Only samples showing detectable responses to the respective positive control were included. Within each assay and Aβ PET status group, stimulation conditions were compared against No Ag condition using paired Wilcoxon signed-rank tests; displayed *P* values were adjusted for multiple testing within each assay using BH FDR. Heatmaps summarize median ICOS+CD25+ responses after pooling participants by Aβ PET status. Abbreviations: CU = cognitively unimpaired older adults; MCI = mild cognitive impairment; OO = oldest-old participants without dementia; EM = effector memory; CM = central memory; AUC = area under the curve; TIV = total intracranial volume; SUVR = standardized uptake value ratio; BH FDR = Benjamini-Hochberg false discovery rate approach; SD = standard deviation.

To determine whether ICOS surface expression on the selected memory T-cell subsets related to longitudinal AD-relevant outcomes, we next tested hippocampal volume change, WMH volume change and episodic memory change. For each outcome, we evaluated three linear regression models separately within CU, MCI and OO participants. Model 1 tested the association between baseline cerebral Aβ load and the longitudinal outcome. Model 2 tested whether each immune-marker feature showed an additive association with the outcome after adjustment for baseline cerebral Aβ load. Model 3 tested whether each immune-marker feature modified the association between baseline cerebral Aβ load and the longitudinal outcome through an Aβ PET × immune-marker interaction (**Extended Data Fig. 4a-c**).

We first focused on hippocampal volume change as a longitudinal marker of neurodegeneration. As expected, higher baseline cerebral Aβ load was associated with more negative hippocampal volume change in CU and MCI participants, consistent with greater hippocampal atrophy (**Extended Data Fig. 4a**, Model 1). In contrast, none of the selected immune-marker features showed additive associations with hippocampal volume change after adjustment for cerebral Aβ load (**Extended Data Fig. 4a**, Model 2). However, interaction models identified significant negative Aβ PET × immune-marker interactions in MCI participants for ICOS signal intensity in CD8 EM and CD8 CM T-cell clusters (**Fig. 4d and Extended Data Fig. 4a**, Model 3). Thus, higher ICOS signal on these CD8 memory T-cell subsets strengthened the negative association between cerebral Aβ load and longitudinal hippocampal volume change, consistent with greater Aβ-associated hippocampal atrophy. Model-based prediction plots illustrated this interaction pattern. In MCI participants with low or mean ICOS signal intensity in CD8 EM or CD8 CM T-cell clusters, predicted hippocampal volume change showed a weaker association with increasing cerebral Aβ load. In contrast, at high ICOS signal intensity, higher Aβ load was associated with more negative predicted hippocampal volume change, indicating stronger Aβ-associated hippocampal atrophy (**Fig. 4e,f**).

Because Aβ pathology and WMH burden can exert clinically relevant co-pathology effects, particularly for cognitive decline,^44^ we next tested whether the selected immune-marker features related to longitudinal WMH accumulation. In contrast to the hippocampal atrophy models, WMH analyses did not identify significant Aβ PET × immune-marker interactions. Instead, ICOS signal on CD8 EM showed an additive association with WMH volume change in CU participants after adjustment for baseline cerebral Aβ load (**Extended Data Fig. 4b**, Model 2). This pattern suggests an Aβ-independent association between CD8 EM T-cell ICOS signal and WMH progression in CU participants.

Finally, we tested whether the selected ICOS immune-marker features related to longitudinal episodic memory change. As expected, higher baseline cerebral Aβ load was associated with more negative episodic memory change in CU and especially MCI participants, indicating Aβ-associated risk of cognitive decline (**Extended Data Fig. 4c**, Model 1). However, the selected immune-marker features did not show additive associations with episodic memory change and did not significantly modify the association between Aβ load and episodic memory change in these models (**Extended Data Fig. 4c**, Models 2 and 3).

### Aβ-reactive, ICOS-centred T-cell responses are detectable in an independent study population 2

To test whether Aβ-associated T-cell findings from study population 1 were reflected in functional antigen responses, we used an independent study population 2 derived from a previously established cohort characterized by cerebral Aβ PET imaging with [11C]-Pittsburgh compound B (PiB-PET).^32^ This population comprised CU participants (n = 14) and participants with MCI (n = 10), with available PBMCs and plasma for *in vitro* antigen-stimulation assays and AD biomarker analyses (**Fig. 4g**, **Table 2 and Extended Data Table 2**). Cerebral Aβ positivity was defined using the same Centiloid threshold as in study population 1. Within this independent population, 10 of 14 CU participants and 6 of 10 MCI participants were PiB-PET Aβ-positive. AD biomarker correlations were consistent with the expected direction of cerebral Aβ pathology: higher Aβ PET signal was associated with lower plasma Aβ42/Aβ40 ratio and higher plasma p-tau181 (**Fig. 4h,i**), establishing study population 2 as an independent, biomarker-characterized sample set.

Study population 2 was then used for an *in vitro* antigen-stimulation workflow in which sorted memory CD4 and CD8 T-cells were CFSE-labelled and co-cultured with autologous monocytes pulsed with peptide pools derived from the N-terminal (Aβ-N), mid-sequence (Aβ-M) and C-terminal (Aβ-C) regions of the Aβ1-42 peptide, or with control antigens (**Fig. 4g**). Proliferating T-cells were identified by flow-cytometric assessment as CFSE^low^ cells. Co-expression of CD25, the IL-2 receptor α-chain and canonical T-cell activation marker, with ICOS was used as an inducible, activation-associated readout within the proliferating memory T-cell compartment. In memory CD4 T-cells from cerebral Aβ-positive participants, all three Aβ peptide pools induced increased frequencies of ICOS+CD25+ cells within the CFSE^low^ compartment compared with unstimulated cultures, with the strongest median response observed after stimulation with the Aβ C-terminal peptide pool (**Fig. 4j**). Influenza antigen stimulation served as a positive control and induced the expected ICOS+CD25+ CD4 T-cell response. In contrast, memory CD4 T-cells from Aβ-negative participants showed no detectable increase in ICOS+CD25+ CFSE^low^ cells after stimulation with any Aβ peptide pool, whereas positive-control stimulation induced a response (**Fig. 4j**). A similar pattern was observed in the memory CD8 T-cell assay. In Aβ-positive participants, pooled Aβ-derived peptides induced increased frequencies of ICOS+CD25+ cells among proliferating CFSE^low^ memory CD8 T-cells, at levels comparable to the EBV positive-control response (**Fig. 4k**). In Aβ-negative participants, pooled Aβ-derived peptides did not induce an ICOS+CD25+ CD8 T-cell response, whereas EBV stimulation induced a weak positive-control response (**Fig. 4k**). Together, these findings show that Aβ peptide-reactive memory CD4 and CD8 T-cell responses with ICOS and CD25 co-expression were detectable in the independent study population 2 and were enriched in Aβ-positive participants.

### Oldest-old participants show altered Aβ pathology-related episodic memory trajectories and resilience-associated immune states

Resilience in AD denotes the ability to maintain cognitive function despite AD pathology.^80^ Because the OO group consisted of individuals of exceptional age with preserved daily-life activities despite a high prevalence of cerebral Aβ pathology,^16^ we asked whether the relationship between cerebral Aβ load and longitudinal episodic memory trajectories differed across CU, MCI and OO participants. In longitudinal mixed-effects models, higher baseline cerebral Aβ load was associated with more negative episodic memory trajectories in CU participants and, more prominently, in MCI participants (**Fig. 5a**). In contrast, OO participants showed an altered relationship of cerebral Aβ load and cognitive change. Although baseline episodic memory performance was lower and longitudinal episodic memory change was more negative in OO groups compared to CU participants (**Extended Data Fig. 1l,m**), modelled trajectories over time as a function of cerebral Aβ load showed a highly variable pattern in OO, rather than the clearer Aβ-associated decline observed in MCI (**Fig. 5a**). Thus, despite advanced age and frequent cerebral Aβ pathology, OO participants showed evidence of altered cognitive vulnerability to Aβ burden, distinct from the MCI pattern. Accordingly, we considered OO+ participants an AD-resilience-enriched group, operationally defined by exceptional age, PET Aβ positivity and absence of dementia, and used this group for subsequent immune analyses.

**Fig. 5:**
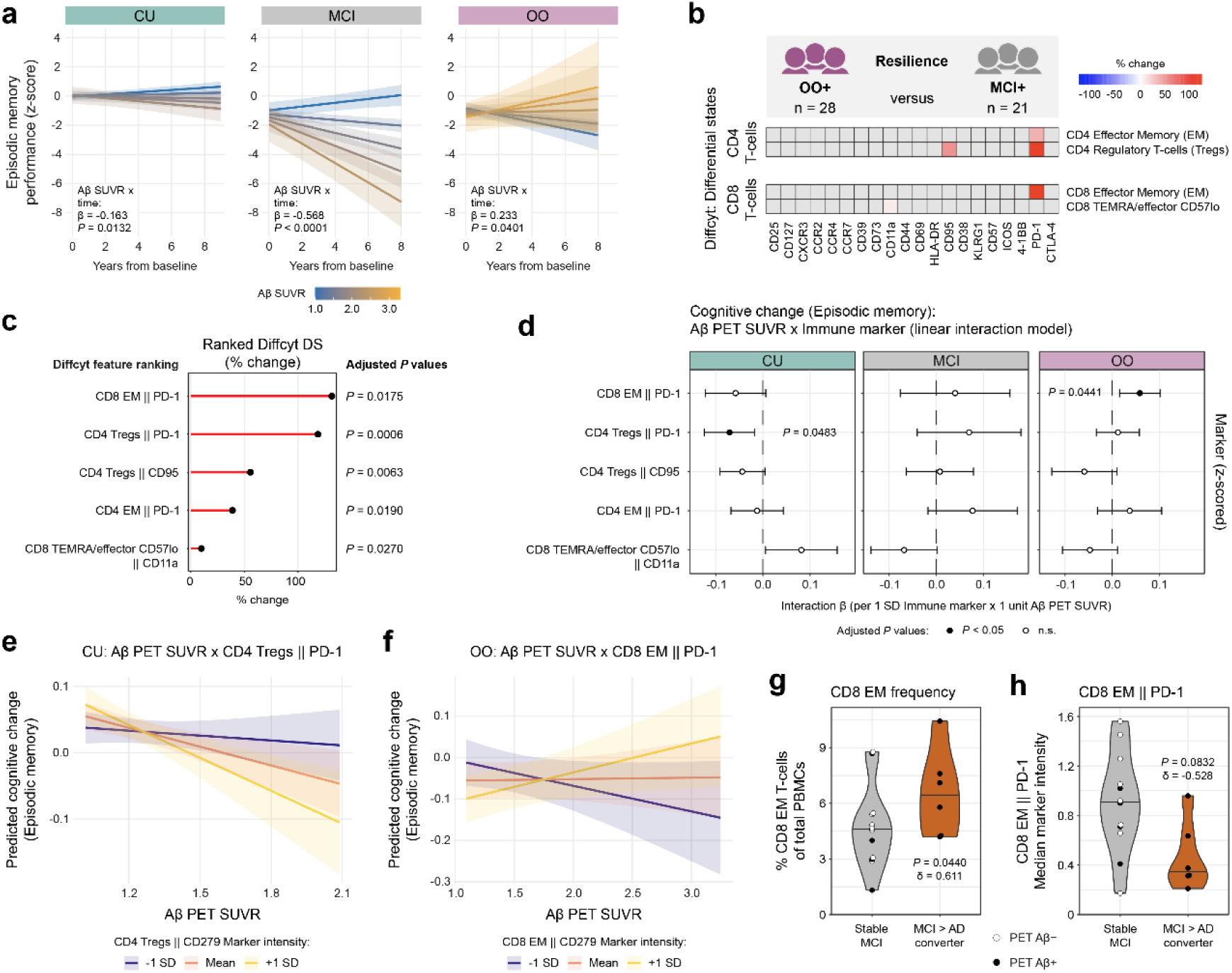
Resilience-associated immune cell states are linked to altered Aβ pathology-related episodic memory trajectories. **a**. Longitudinal mixed-effects models of predicted episodic memory performance over follow-up time since baseline as a function of cerebral Aβ load in CU, MCI and OO. Models were fitted separately within each diagnostic group and included time from baseline, Aβ PET SUVR and their interaction, with adjustment for baseline age, sex and years of education. Lines show model-estimated episodic memory trajectories across Aβ SUVR levels, with shaded areas indicating 95% confidence intervals. Aβ SUVR scale is standardized across diagnostic groups. β values indicate the time × Aβ SUVR interaction term; *P* values were adjusted globally using BH FDR. **b**. Diffcyt differential state analysis of the resilience contrast comparing Aβ-positive OO with Aβ-positive MCI (OO+ versus MCI+). Analyses were adjusted for sex and CMV IgG. To reduce confounding by age-associated immune differences, significant features overlapping with the OO− versus CU− age-control contrast were removed. Heatmap colours indicate raw percentage change in marker expression. Grey tiles indicate features not passing the BH FDR-adjusted threshold of *P* < 0.05; landmark clusters without significant hits are not shown. **c**. Ranking of resilience-associated Diffcyt differential state results. Features were ranked by absolute raw percentage change. Line colour indicates the direction of change, as in b. Labels indicate BH FDR-adjusted *P* values. **d**. Linear interaction models testing whether resilience-associated immune-marker expression modified the association between cerebral Aβ load and episodic memory change. Models included baseline age, sex, years of education, Aβ PET SUVR, z-scored immune-marker expression and their interaction. Forest plots show the Aβ SUVR × immune-marker interaction β estimate and 95% confidence interval; β reflects the change in Aβ PET-episodic memory change association per 1 SD higher immune-marker expression. Filled circles indicate BH FDR-adjusted *P* < 0.05 within each diagnostic group. **e,f**. Model-based prediction plots for significant interactions. Predicted episodic memory change is shown across the observed range of Aβ PET SUVR at low (−1 SD), mean and high (+1 SD) immune-marker expression for CD4 Tregs || PD-1 in CU (e) and CD8 EM || PD-1 in OO (f), illustrating how immune-marker levels modify the association between cerebral Aβ load and cognitive change. Predictions represent conditional estimates from an unconstrained linear model. **g,h**. Baseline CD8 EM T-cell characteristics in longitudinally stable MCI participants and participants who subsequently converted from MCI to AD. Violin plots show frequency of CD8 EM T-cells (g) and PD-1 marker expression on CD8 EM T-cells (h). Black filled dots represent PET Aβ+ subjects at baseline. Group differences were assessed using two-sided Wilcoxon rank-sum tests with adjustment for sex and CMV IgG status. Cliff’s delta is shown as non-parametric effect-size estimate. Abbreviations: CU = cognitively unimpaired older adults; MCI = mild cognitive impairment; OO = oldest-old participants without dementia; SUVR = standardized uptake value ratio; BH FDR = Benjamini-Hochberg false discovery rate approach; SD = standard deviation; EM = effector memory; Tregs = regulatory T-cells; PD-1 = programmed cell death protein 1.

To identify immune-cell states associated with this resilience-enriched phenotype, we next applied diffcyt differential-state analysis to the resilience contrast comparing OO+ with MCI+ participants. This contrast compared two Aβ-positive groups with divergent clinical vulnerability: OO participants without dementia and MCI participants with AD pathology. Models were adjusted for sex and CMV IgG titre. Because OO participants were much older than CU and MCI participants, we additionally removed significant features that overlapped with the OO− versus CU− age-control contrast, reducing the contribution of age-associated immune differences. The resilience contrast identified a distinct T-cell state pattern characterized by higher PD-1 signal in several T-cell subsets, including CD8 EM, CD4 EM and CD4 Tregs, together with higher CD95 signal in CD4 Tregs and higher CD11a signal in CD8 TEMRA/effector CD57lo T-cells (**Fig. 5b**). Ranking features by raw percentage change identified CD8 EM || PD-1 as the strongest resilience-associated differential-state feature (**Fig. 5c**).

Next, we applied the longitudinal modelling framework, used earlier for ICOS immune-marker features, to the resilience-associated immune-marker features with episodic memory change as outcome. Model 2 analyses did not identify additive associations between resilience-associated immune-marker features and episodic memory change after adjustment for baseline cerebral Aβ load (**Extended Data Fig. 4d**, Model 2). In contrast, Model 3 interaction analyses identified opposing Aβ PET × immune-marker interactions involving PD-1 signal in T-cell subsets: a negative interaction for CD4 Tregs || PD-1 in CU participants and a positive interaction for CD8 EM || PD-1 in OO participants (**Fig. 5d and Extended Data Fig. 4d**, Model 3). Model-based prediction plots clarified the direction of these interactions in detail. In CU, high baseline PD-1 signal in CD4 Tregs was associated with a more negative relationship between cerebral Aβ load and predicted episodic memory change, indicating stronger Aβ-associated cognitive decline (**Fig. 5e**). In contrast, in OO, high baseline PD-1 signal in CD8 EM T-cells was associated with an attenuated relationship between cerebral Aβ load and episodic memory change, with predicted episodic memory change remaining relatively preserved across increasing cerebral Aβ load (**Fig. 5f**). Thus, CD8 EM || PD-1 showed a resilience-linked interaction pattern in OO participants, compatible with weaker Aβ-associated cognitive vulnerability at higher PD-1 signal.

Finally, we explored whether baseline immune characteristics were linked to subsequent longitudinal diagnostic stability or conversion. During follow-up, six MCI participants converted to AD (**Extended Data Table 3**). We therefore compared baseline immune features between longitudinally stable MCI participants and MCI-to-AD converters. Although CD8 EM T-cell frequencies were not significantly altered in the cross-sectional CU−, CU+ and MCI+ Aβ-pathology comparison (**Fig. 2d**), they were higher at baseline in MCI-to-AD converters than in stable MCI participants (**Fig. 5g**). Conversely, PD-1 signal intensity within CD8 EM T-cells was lower in MCI-to-AD converters than in stable MCI participants (**Fig. 5h**), complementing the OO interaction analysis in which higher CD8 EM PD-1 signal was linked to attenuated Aβ-associated episodic memory decline.

## Discussion

Across the AD continuum assessed here, our findings suggest two separable adaptive immune patterns: an ICOS-centred activation state in CD4 and CD8 memory T-cells associated with cerebral Aβ pathology and, in MCI, greater Aβ-associated hippocampal atrophy; and a PD-1-centred CD8 EM T-cell state linked to a resilience-associated clinical profile. Aβ-derived peptides induced proliferative ICOS+CD25+ memory CD4 and CD8 T-cell responses predominantly in Aβ-positive participants in an independent study population, linking the ICOS phenotype to Aβ-directed immune reactivity. Conversely, higher PD-1 expression on CD8 EM T-cells was associated with an attenuated relationship between cerebral Aβ load and episodic-memory decline in OO participants, and distinguished longitudinally stable MCI participants from MCI-to-AD converters. These findings identify co-stimulatory and co-inhibitory T-cell states as distinct correlates of vulnerability and resilience, without establishing causal effects on neurodegeneration.

The clinical and biomarker architecture of the cohort provides an important context for distinguishing AD-related changes from processes accompanying advanced age. In the MCI group, higher cerebral Aβ load and increased plasma p-tau217 correlated with hippocampal atrophy, WMH accumulation and episodic-memory decline, consistent with established relationships between Aβ pathology, neurodegeneration and clinical progression.^44,71,72,81^ In contrast, both Aβ-positive and Aβ-negative OO groups showed higher NfL and GFAP plasma levels, greater hippocampal atrophy and WMH accumulation, and worse episodic-memory performance, irrespective of Aβ status. This pattern is compatible with increasingly heterogeneous neuroaxonal, astroglial and vascular injury at advanced age, as well as age-related brain atrophy that may contribute to reduced cognitive performance independently of AD.^82^ Peripheral NfL rises with age and is associated with structural brain changes,^83,84^ and both plasma NfL and GFAP have been shown to be elevated in another oldest-old cohort comprising individuals aged ≥90 years with preserved cognitive function.^85^ The relatively sparse biomarker correlations in OO do not imply an absence of biologically relevant brain changes. The limited number of within-group associations surviving FDR correction may instead reflect the restricted clinical range imposed by selection for absence of dementia and preserved everyday function, together with increasing biological heterogeneity, co-pathology and survivor selection at very advanced ages. The latter may have altered biomarker-outcome coupling if *APOE* ε4-associated trajectories led to cognitive impairment or death before recruitment. Consistent with this, the proportion of *APOE* ε4 carriers was 12.5% in our OO group compared to 24.6% in combined CU and MCI groups, and only 3% in the cohort of individuals aged ≥90 years with preserved cognitive function studied by Gómez-Tortosa and colleagues.^85^ Moreover, the episodic-memory composite was standardized against the baseline CU distribution rather than an age-specific OO reference, making longitudinal trajectories and their relationship to cerebral Aβ load more informative than baseline differences alone. Previous work indicates that hippocampal structure becomes more closely related to memory at older ages,^74^ while the association between Aβ and memory may be overshadowed by atrophy and additional age-related pathologies in later life.^86^ Together, these observations support interpreting the OO group as a heterogeneous, resilience-enriched population in which cerebral Aβ load is less tightly coupled to clinical outcomes, and not as a group entirely lacking age-related brain injury or potential consequences of Aβ pathology.

Age-associated immune remodelling provided an important background for interpreting the Aβ pathology-associated changes. Consistent with established features of immunosenescence and inflammaging,^87,88^ the healthy-aging comparison showed contraction of naïve CD4 and CD8 T-cell compartments and broader shifts in peripheral immune composition. Persistent antigenic exposure, particularly to CMV, is an important contributor to such remodelling in later life.^89–91^ In our cohort, CMV IgG titre was the serological measure most strongly associated with differentiated T-cell frequencies, and subsequent analyses were therefore adjusted for CMV IgG titre. Within the AD continuum, CD8 TEMRA/effector T-cells were more frequent in Aβ-positive CU and MCI participants, recapitulating our previous observations across peripheral blood and CSF.^32,33^ Increased and clonally expanded CD8 TEMRA cells have likewise been reported in AD and ALS,^30,92^ indicating that this compositional change might not be unique to AD pathology.

Cell frequency and surface marker expression within phenotypically defined subsets provide complementary information: the former captures changes in compartment size, whereas the latter can reveal activation-, differentiation-, or co-signalling-associated phenotypes within otherwise similar cellular populations. ICOS is an inducible CD28-family co-stimulatory receptor whose engagement by ICOSL can support T-cell proliferation, survival, differentiation and cytokine production, whereas its functional consequences depend on the cellular and inflammatory context.^40,93^ Here, ICOS was the most recurrent Aβ-associated marker across CD4 and CD8 memory T-cell compartments. Notably, ICOS was also increased on CD8 TEMRA/effector cells in the MCI+ versus CU− and MCI+ versus MCI− contrasts, linking the CD8 TEMRA/effector frequency finding to a potentially altered co-signalling phenotype. However, ICOS on CD8 TEMRA/effector cells did not emerge from the continuous cerebral Aβ load screen and RF-RFE overlap, suggesting a stage- or context-dependent association. The more robust Aβ pathology-associated signature comprised ICOS on CD4 EM, CD8 EM and CD8 CM T-cells. Recurrence of the same surface receptor across several memory T-cell compartments supports a shared molecular axis more strongly than an isolated cluster difference, while ICOS abundance alone establishes neither antigen specificity nor a defined effector function. In OO participants, the elevated ICOS marker intensity background already present in the Aβ-negative group is compatible with the concept of inflammaging and advanced-age immune remodelling, and likely reduced the separation of ICOS levels according to Aβ status. The ROC analyses mirrored this pattern: adding ICOS expression on CD4 EM, CD8 EM and CD8 CM T-cells to a reference model yielded high discrimination of Aβ status in CU and MCI participants, with AUCs of 0.887 and 0.982, respectively. The immune model significantly outperformed the plasma-biomarker model in CU and showed weaker evidence for improvement in MCI participants. In OO, by contrast, the immune model showed lower discrimination than the plasma-biomarker model. Thus, the ROC results support the consistency of the ICOS signature in CU and MCI, whereas its association with Aβ status appears to be attenuated against the altered immune background of advanced age. Validation in independent cohorts will be required to establish its potential biomarker value.

Functional support for the ICOS-centred findings came from antigen-stimulation experiments in the independent study population 2. Aβ1-42-derived peptide pools induced proliferative ICOS+CD25+ memory CD4 and CD8 T-cell responses predominantly in PET Aβ-positive participants. In memory CD4 T-cells, peptide pools spanning the N-terminal, mid-sequence and C-terminal Aβ1-42 regions all elicited responses, with the strongest response observed for the C-terminal pool. In memory CD8 T-cells, a combined Aβ peptide pool elicited the ICOS+CD25+ response. These findings extend previous evidence of Aβ-reactive T-cells in older individuals and patients with AD^38,39^ and link the Aβ pathology-associated ICOS phenotype to the capacity to mount an Aβ-peptide-directed response. However, these experiments do not constitute a direct replication of the *ex vivo* CyTOF findings, as the functional assays measured ICOS and CD25 among sorted memory T-cells that had proliferated during six days of co-culture with peptide-pulsed autologous monocytes. The enrichment of responses in PET Aβ-positive participants suggests previous priming or expansion of Aβ-reactive memory T-cells, but does not establish the origin, duration or timing of antigenic exposure, nor the kinetics of ICOS induction, which was assessed only on day 6. These findings relate to our previous work using a 16-hour PBMC assay, which identified Aβ-responsive memory CD4 T-cells through OX40 and 4-1BB expression, with the strongest responses elicited by mid-sequence and C-terminal Aβ1-42 pools in Aβ-positive cognitively unimpaired participants, but broadly reduced responses in MCI+ participants.^33^ The two assays captured different response phases: short-term activation-marker induction versus the phenotype of cells that had survived and proliferated over six days. Together with exhaustion-like CD4 and CD8 T-cell phenotypes reported in Aβ-positive MCI,^94^ the previous OX40- and 4-1BB-associated findings support MCI-associated T-cell hyporesponsiveness. However, the present *ex vivo* and stimulation data indicate that this hyporesponsiveness is not uniform across co-stimulatory pathways, as ICOS-centred signals were increased in both Aβ-positive CU and MCI participants. This distinction is biologically plausible because these co-stimulatory pathways are non-redundant: OX40 and 4-1BB are TNF receptor-superfamily members signalling through TRAF-dependent pathways, whereas ICOS belongs to the CD28 family and signals prominently through PI3K.^41^ In addition, inter-individual HLA haplotype diversity among study participants may have shaped peptide presentation by autologous monocytes, thereby contributing to variability in detectable Aβ-reactive T-cell responses and subgroup-level response patterns.

Revisiting whether Aβ-associated immune responses are beneficial or detrimental during disease progression, we found that the ICOS-centred phenotype was most clearly linked to structural vulnerability in MCI. ICOS expression on CD8 EM and CM T-cells showed no Aβ-independent association with hippocampal volume change; however, both immune-marker features interacted with baseline cerebral Aβ load, such that higher ICOS expression strengthened the association between cerebral Aβ and hippocampal atrophy in MCI. Comparable Aβ PET × ICOS interactions were not detected for WMH progression or episodic-memory decline, although cerebral Aβ load itself was associated with episodic-memory decline. This dissociation is compatible with hippocampal atrophy becoming detectable before a corresponding, potentially ICOS-associated, cognitive effect, without establishing this temporal sequence.^72,95^ The broader literature provides plausible, but currently unproven, mechanistic context for CD8 T-cell involvement in neurodegeneration. In Aβ-pathology mouse models, plaque-associated CD8 T-cells accumulated with progressing amyloidosis.^96^ In a 3D human neuroimmune AD model, CXCL10-CXCR3-dependent CD8 T-cell infiltration increased microglial activation, neuroinflammation and neuronal damage, whereas blockade of this axis attenuated T-cell infiltration and neurodegeneration.^97^ A downstream or co-pathology route involving tau is also plausible: extravascular T-cells in human AD brain were predominantly CD8 T-cells and correlated with tau rather than Aβ pathology,^29^ while a tauopathy mouse model linked activated, clonally expanded CD8 T-cells to a microglial inflammatory circuit in which combined CD4/CD8 T-cell depletion or IFNγ blockade attenuated brain atrophy.^34^ Together, these studies support the possibility that higher ICOS expression on peripheral CD8 memory T-cells marks an immune context in which Aβ-associated neurodegeneration is greater, potentially involving T-cell trafficking, microglia-modifying effects or interactions with downstream tau pathology. However, peripheral ICOS expression establishes neither CNS entry nor cytotoxic or microglia-modifying activity; ICOS-ICOSL axis perturbation combined with assessment of T-cell trafficking and CNS effector function will be required to distinguish marker from mediator.

Previous work in the OO cohort linked cognitive reserve and current physical activity to preserved cognition and lower cerebral Aβ load, respectively.^16^ Genetic and human brain single-cell studies also implicate immune-response pathways and distinct microglial trajectories in resilience to Aβ pathology at advanced age.^22–24^ In our data, OO+ participants showed higher PD-1 signal across several T-cell subsets compared to MCI+ participants, with PD-1 on CD8 EM T-cells emerging as the strongest resilience-associated feature. Higher CD8 EM PD-1 did not show a direct association with episodic-memory change but attenuated the relationship between cerebral Aβ load and episodic-memory decline in OO. Likewise, higher CD8 EM PD-1 in stable MCI participants compared to MCI-to-AD converters provided an exploratory observation in the same direction. Because PD-1 is an activation-inducible co-inhibitory receptor that constrains TCR and CD28 signalling after PD-L1 ligand engagement,^42^ these findings are consistent with a more restrained CD8 EM phenotype. This is in line with evidence that CXCR6+PD-1+ brain-resident CD8 T-cells can limit pro-inflammatory microglial responses, Aβ deposition and cognitive decline in Aβ-pathology mouse models.^35^ However, PD-1 associations were subset- and context-dependent: in CU participants, higher PD-1 signal on CD4 Tregs was associated with stronger Aβ-related episodic-memory decline. This extends our previous cross-sectional observation of a less immunosuppressive CD4 Treg transcriptome in preclinical AD^33^ by linking altered Treg biology to longitudinal cognitive vulnerability rather than only to potentially permissive Aβ-reactive immunity. This interpretation is compatible with insufficient regulatory control, given the evidence that PD-1 signalling can restrain Treg activation and suppressive capacity.^98^ Surface PD-1 expression alone, however, establishes neither ligand engagement, downstream signalling nor impaired suppressive function. Intervention studies further indicate that PD-1-PD-L1 biology is context-dependent. PD-1-PD-L1 blockade reduced pathology through systemic immune activation and recruitment of monocyte-derived macrophages in Aβ- and tau-pathology mouse models,^99,100^ whereas another study reported no effect of PD-1 immunotherapy on Aβ pathology.^101^ Blockade of the PD-1-PD-L1 axis in tauopathy models resulted not only in mobilization of monocytes to the brain and attenuation of atrophy, but also in additional recruitment of CD4 Tregs.^34^ This is conceptually consistent with our observation that PD-1 expression on Tregs was associated with Aβ-dependent cognitive decline, raising the possibility that PD-1 blockade could alter this axis. In early symptomatic AD, short-lived anti-PD-L1 treatment induced a transient, dose-dependent increase in peripheral PD-1+ICOS+ CD4 memory T-cells, but the phase 1b study neither established whether these cells were beneficial or detrimental nor demonstrated clinical efficacy.^102^ Given our association of ICOS-expressing memory T-cells with Aβ pathology and Aβ-associated hippocampal atrophy, induced PD-1+ICOS+ T-cell states warrant cell-resolved longitudinal monitoring alongside functional and clinical outcomes.

A major strength of this study is the integration of high-dimensional immune profiling across a biomarker-defined AD and aging continuum, spanning CU+ participants as potential preclinical AD,^3,4^ MCI+ participants as prodromal AD, and exceptionally old individuals without dementia with or without cerebral Aβ pathology, including an OO+ resilience-enriched phenotype. Immune features were related to cross-sectional fluid and imaging biomarkers and to longitudinal cognitive, hippocampal and vascular outcomes over up to eight years, and were complemented by Aβ-peptide stimulation experiments in an independent study population. Limitations include the single-centre design and modest diagnostic subgroup sizes, which restrict generalizability and make interaction and converter analyses exploratory. Tau PET and fluid biomarkers specific for later-stage tau aggregation were unavailable; unmeasured tau pathology may therefore have contributed to the imaging and cognitive outcomes.

Taken together, our findings favour a model in which insufficiently restrained adaptive immune responses may accompany or contribute to vulnerability in early AD, whereas selective restraint of potentially cytotoxic CD8 memory T-cells may support resilience. The divergent ICOS- and PD-1-centred associations indicate that adaptive immunity in early AD is not uniformly harmful or protective, but depends on cell type, activation state, antigen context and disease stage. These findings caution against indiscriminate global immune activation approaches and support precision immunomodulation informed by the same parameters. Co-stimulatory ICOS may provide an accessible readout and experimental therapeutic target, whereas antigen-specific immunosuppression, by CAR-Tregs for example, could offer a complementary tolerance-induction strategy.^103^ Longitudinal tracking of co-stimulatory immune states and antigen-specific clonal TCR expansions, enabled by scalable TCR-antigen mapping approaches,^60,104^ may advance immune-related AD biomarker strategies and precision immunomodulation.

## Extended Data

**Extended Data Table 1: Characteristics and available data across study participant subgroups in study population 1.**

Summary of demographic, genetic, neuropsychological, imaging, plasma biomarker and inflammatory/infection-related variables across study groups. The Excel workbook contains two sheets: 1a_Characteristics, showing cohort summaries and statistical test results; and 1b_Available n, showing the number of available observations per variable and group.

For normal distributed variables summarized as mean ± SD, global group differences were assessed using one-way ANOVA, and pairwise comparisons against reference group CU− were performed using Welch’s t-tests. For non-normal distributed variables summarized as median with [IQR], global group differences were assessed using non-parametric Kruskal-Wallis tests, and pairwise comparisons against CU− were performed using Dunn’s tests. For categorical variables, global group differences were assessed using χ² tests when all expected contingency-table cell counts were ≥ 5; Fisher’s exact test was used instead when at least one expected cell count was < 5 or when χ² testing was not appropriate. Pairwise categorical comparisons against CU− were performed using Fisher’s exact tests. The test statistics column reports the global test across groups. Symbols shown within individual group cells indicate pairwise comparisons of that group against CU− after BH FDR correction within the corresponding variable: ^+^ *P* < 0.1; * *P* < 0.05; ** *P* < 0.01; *** *P* < 0.001; **** *P* < 0.0001. Absence of a symbol indicates that the pairwise comparison against CU− was not significant after row-wise BH FDR correction. Except for baseline age and subject counts, values for the young control group were not available and are marked as “na”. CDR-SB was not available for OO− and OO+ and is therefore marked as “na”.

Abbreviations: Aβ = amyloid-β; APOE = apolipoprotein E; AE/ml = assay-reported units per ml; BH FDR = Benjamini-Hochberg false discovery rate approach; CDR-SB = Clinical Dementia Rating Sum of Boxes; CMV = cytomegalovirus; CRP = C-reactive protein; CU = cognitively unimpaired older adults; FMM = [18F]-flutemetamol; GFAP = glial fibrillary acidic protein; IQR = interquartile range; MCI = mild cognitive impairment; MRI = magnetic resonance imaging; NfL = neurofilament light; NP = neuropsychological; OO = oldest-old participants without dementia; PET = positron emission tomography; p-tau217 = phosphorylated tau 217; SD = standard deviation; SUVR = standardized uptake value ratio; TIV = total intracranial volume; WMH = white matter hyperintensity.

**Extended Data Table 2: Characteristics and available data across study participant subgroups in study population 2.**

Summary of demographic, genetic, neuropsychological, imaging, plasma biomarker and inflammatory/infection-related variables across study groups. The Excel workbook contains two sheets: 2a_Characteristics, showing cohort summaries and statistical test results; and 2b_Available n, showing the number of available observations per variable and group.

For normal distributed variables summarized as mean ± SD, global group differences were assessed using one-way ANOVA, and pairwise comparisons against reference group CU− were performed using Welch’s t-tests. For non-normal distributed variables summarized as median with [IQR], global group differences were assessed using non-parametric Kruskal-Wallis tests, and pairwise comparisons against CU− were performed using Dunn’s tests. For categorical variables, global group differences were assessed using Fisher’s exact test. The test statistics column reports the global test across groups. Symbols shown within individual group cells indicate pairwise comparisons of that group against CU− after BH FDR correction within the corresponding variable: ^+^ *P* < 0.1; * *P* < 0.05; ** *P* < 0.01; *** *P* < 0.001; **** *P* < 0.0001. Absence of a symbol indicates that the pairwise comparison against CU−was not significant after row-wise BH FDR correction.

Abbreviations: Aβ = amyloid-β; APOE = apolipoprotein E; BH FDR = Benjamini-Hochberg false discovery rate approach; CRP = C-reactive protein; CU = cognitively unimpaired older adults; IQR = interquartile range; MCI = mild cognitive impairment; PET = positron emission tomography; PiB = [11C]-Pittsburgh Compound-B; p-tau181 = phosphorylated tau 181; SD = standard deviation.

**Extended Data Table 3: Longitudinal diagnostic classification and conversion patterns in study population 1.**

Summary of availability of valid follow-up diagnosis and classification as converters, transient/non-persistent diagnostic changes, stable CU/MCI/OO, or other non-stable followed participants. Converter counts are additionally shown by specific diagnostic transition.

CU = cognitively unimpaired older adults; MCI = mild cognitive impairment; OO = oldest-old participants without dementia; PRADL = probable Alzheimer’s disease late-onset.

## Acknowledgements

We would like to thank all the volunteers who kindly participated in the studies. We thank all study physicians and neuropsychologists for their contribution to the assessments and conduct of the study. Samples from participants were collected at the Center for Prevention and Dementia Therapy (Institute for Regenerative Medicine, University of Zurich) by study nurses led by Esmeralda Gruber. We would like to thank the Cytometry Facility (University of Zurich) for technical assistance, and the Institute of Medical Virology (University of Zurich) for virus serology analysis. Finally, we would like to thank Hayder Shweliyya and his team at the Sahlgrenska University Hospital for fluid biomarker analyses.

## Author contributions

Conceptualization, A.G., V.T., and C.G.; Methodology, A.M., D.B., C.R., M.K., T.K., and H.Z.; Software, A.M., D.B., and C.G.; Formal Analysis, A.M., D.B., and C.G.; Investigation, A.M., D.B., and C.G.; Resources, C.H., A.G., and V.T.; Data Curation, A.M., D.B., and C.G.; Writing - Original Draft, C.G.; Writing - Review & Editing, A.M., D.B., H.Z., M.T.F., L.K., F.S., A.G., V.T., and C.G.; Visualization, A.M., D.B., C.R., and C.G; Supervision and Project Administration, A.G., V.T., and C.G.; Funding Acquisition, C.R., H.Z., C.H., R.M.N., A.G., and C.G.

## Consent statement

All study participants gave written informed consent. All studies and further use for the current analyses were approved by the local ethics committee (Kantonale Ethikkommission Zürich) and conducted in accordance with their guidelines and the Declaration of Helsinki.

## Supplementary Information

**Supplementary Table 1:** Heavy metal-labeled antibodies used for mass cytometry.

| Isotope | Metal | Antigen | Clone | Antibody Supplier | Source/ Identifier | Category |
| --- | --- | --- | --- | --- | --- | --- |
| 89 | Y | CD45 | HI30 | Fluidigm | Cat# 3089003B,<br>RRID: AB_2938863 | barcoding |
| 106 | Cd | CD45 | HI30 | BioLegend | in-house labeling | barcoding |
| 110 | Cd | CD45 | HI30 | BioLegend | in-house labeling | barcoding |
| 111 | Cd | CD45 | HI30 | BioLegend | in-house labeling | barcoding |
| 112 | Cd | CD45 | HI30 | BioLegend | in-house labeling | barcoding |
| 113 | In | CD45 | HI30 | BioLegend | in-house labeling | barcoding |
| 114 | Cd | CD45 | HI30 | BioLegend | in-house labeling | barcoding |
| 116 | Cd | CD8a | IA6-2 | Fluidigm | in-house labeling | surface |
| 141 | Pr | CD45RA | HI100 | BioLegend | in-house labeling | surface |
| 142 | Nd | CD19 | HIB19 | Fluidigm | Cat# 3142001,<br>RRID: AB_2651155 | surface |
| 143 | Nd | CD123 | 6H6 | Fluidigm | Cat# 3143014B,<br>RRID: AB_2811081 | surface |
| 144 | Nd | CD69 | FN50 | Fluidigm | Cat# 3144018,<br>RRID: AB_2687849 | surface |
| 145 | Nd | CD4 | RPA-T4 | Fluidigm | Cat# 3145001B,<br>RRID: AB_2661789 | surface |
| 146 | Nd | CD11a | HI111 | BioLegend | in-house labeling | surface |
| 147 | Sm | CD303<br>(BDCA2) | 201A | Fluidigm | Cat# 3147009B,<br>RRID: AB_2714153 | surface |
| 148 | Nd | CD137 | 4B4-1 | BioLegend | in-house labeling | surface |
| 149 | Sm | CD25 | 2A3 | Fluidigm | Cat# 3149010B,<br>RRID: AB_2756416 | surface |
| 150 | Nd | CD127 | A019D5 | BioLegend | in-house labeling | surface |
| 151 | Eu | CD14 | M5E2 | Fluidigm | Cat# 3151009B,<br>RRID: AB_2810244 | surface |
| 152 | Sm | KLRG1 | 13F12F2 | ThermoFisher | in-house labeling | surface |
| 153 | Eu | CD192 | K036C2 | Fluidigm | Cat# 3153023B,<br>RRID: N/A | surface |
| 154 | Sm | CD1c<br>(BDCA1) | AD5-8E7 | Miltenyi<br>Biotec | in-house labeling | surface |
| 155 | Gd | CD279<br>(PD-1) | EH12.2H7 | Fluidigm | Cat# 3155009B,<br>RRID: AB_2687854 | surface |
| 156 | Gd | CD183 | G025H7 | Fluidigm | Cat# 3156004B,<br>RRID: N/A | surface |
| 158 | Gd | CD194 | L291H4 | Fluidigm | Cat# 3158032A,<br>RRID: AB_2893003 | surface |
| 159 | Tb | CD197 | G043H7 | Fluidigm | Cat# 3159003,<br>RRID: AB_2714155 | surface |
| 160 | Gd | CD28 | CD28.2 | Fluidigm | Cat# 3160003B,<br>RRID: AB_2868400 | surface |
| 161 | Dy | CD152 | 14D3 | Fluidigm | Cat# 3161004B,<br>RRID: N/A | surface |
| 162 | Dy | TCRgd | B1 | BioLegend | in-house labeling | surface |
| 163 | Dy | CD57 | HCD57 | Fluidigm | Cat# 3163022B,<br>RRID:AB_2756434 | surface |
| 164 | Dy | CD95 | DX2 | Fluidigm | Cat# 3164008B,<br>RRID: N/A | surface |
| 165 | Ho | CD39 | A1 | BioLegend | in-house labeling | surface |
| 166 | Er | CD44 | BJ18 | Fluidigm | Cat# 3166001B,<br>RRID: AB_2744692 | surface |
| 167 | Er | CD27 | L128 | Fluidigm | Cat# 3167006B,<br>RRID: AB_2811093 | surface |
| 168 | Er | CD73 | AD2 | Fluidigm | Cat# 3168015B,<br>RRID: AB_2810249 | surface |
| 169 | Tm | CD45 | HI30 | BioLegend | in-house labeling | barcoding |
| 170 | Er | CD3 | UCHT1 | Fluidigm | Cat# 3170001,<br>RRID: AB_2661807 | surface |
| 171 | Yb | CD45 | HI30 | BioLegend | in-house labeling | barcoding |
| 172 | Yb | CD38 | HIT2 | Fluidigm | Cat# 3172007B,<br>RRID: AB_2756288 | surface |
| 173 | Yb | CD45 | HI30 | BioLegend | in-house labeling | barcoding |
| 174 | Yb | HLA-DR | L243 | Fluidigm | Cat# 3174001,<br>RRID: AB_2665397 | surface |
| 175 | Lu | CD278<br>(ICOS) | C398.4A | Fluidigm | Cat# 3175039B,<br>RRID: AB_2905647 | surface |
| 176 | Yb | CD56 | NCAM16.2 | Fluidigm | Cat# 3176008,<br>RRID: AB_2661813 | surface |
| 209 | Bi | CD16 | 3G8 | Fluidigm | Cat# 3209002B,<br>RRID: AB_2756431 | surface |

**Supplementary Table 2:**
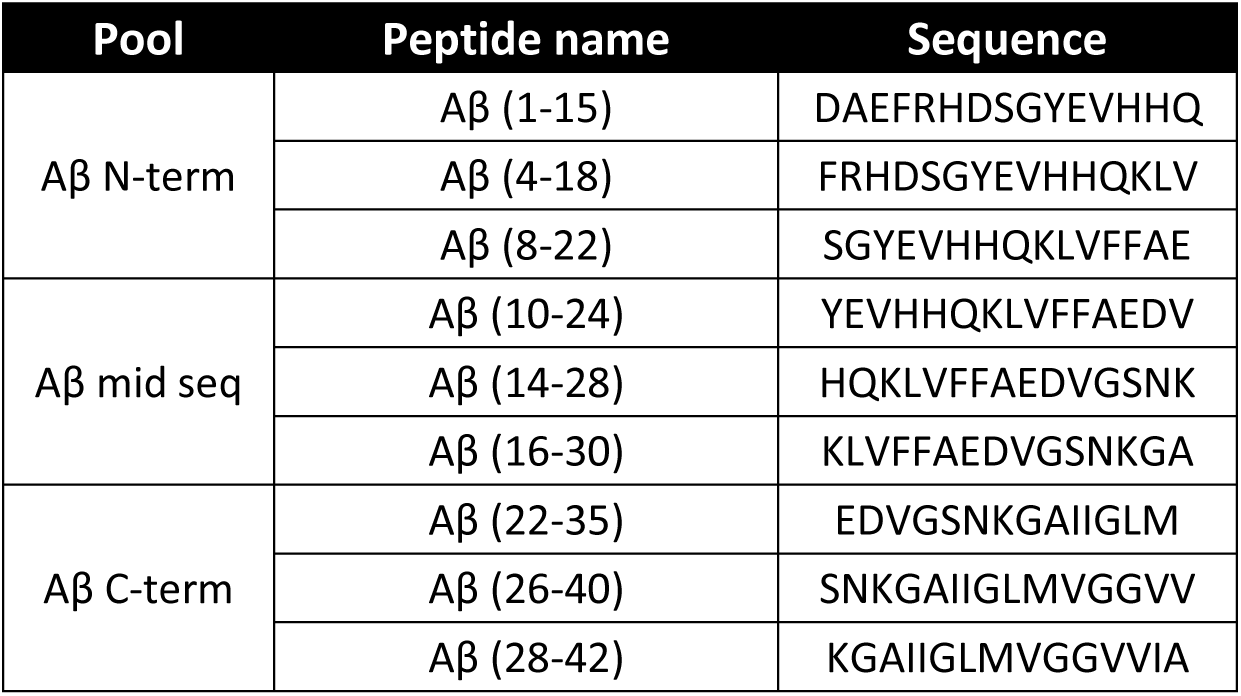
Linear Aβ1-42-derived epitopes used for T-cell antigen presentation. Peptide epitopes were synthesized as 15-mers and are described here with amino acid sequence and position in the full-length peptide of origin.

**Supplementary Table 3:** Fluorochrome-labeled antibody panel used for T-cell antigen-stimulation assays.

| Fluorochrome | Antigen | Clone | Supplier<br>(Cat. No.) | Dilution | Category |
| --- | --- | --- | --- | --- | --- |
| Live/Dead Aqua | Viability dye | – | Invitrogen (L34961) | 1:1000 | L/D |
| BV785 | CD3 | UCHT1 | BioLegend (300472) | 1:100 | Surface (sorting) |
| PE/Dazzle 594 | CD4 | RPA-T4 | BioLegend (300548) | 1:500 | Surface (sorting) |
| APC-Fire 750 | CD8 | RPA-T8 | BioLegend (301066) | 1:80 | Surface (sorting) |
| BV650 | CD45RA | HI100 | BioLegend (304136) | 1:500 | Surface (sorting) |
| BV421 | CCR7 | G043H7 | BioLegend (353208) | 1:80 | Surface (sorting) |
| PE-Cy5 | CD14 | RMO52 | Beckman Coulter | 1:30 | Surface (sorting) |
| FITC | CD19 | HIB19 | BD Biosciences (555412) | 1:20 | Surface (sorting) |
| PE-Cy5 | CD56 | N901 | Beckman Coulter (A07789) | 1:30 | Surface (sorting) |
| PE-Cy5 | CD25 | B1.49.9 | Beckman Coulter (IM2646) | 1:30 | Surface (sorting) |
| PE | CD25 | M-A251 | BD Biosciences (555432) | 1:20 | Activation (day 6) |
| Pacific Blue | ICOS | H4A3 | BioLegend (313522) | 1:100 | Activation (day 6) |
| CFSE | CellTrace™ CFSE | – | Invitrogen | 0.5 µM | Proliferation tracking |

---

**Extended Data Fig. 1:**
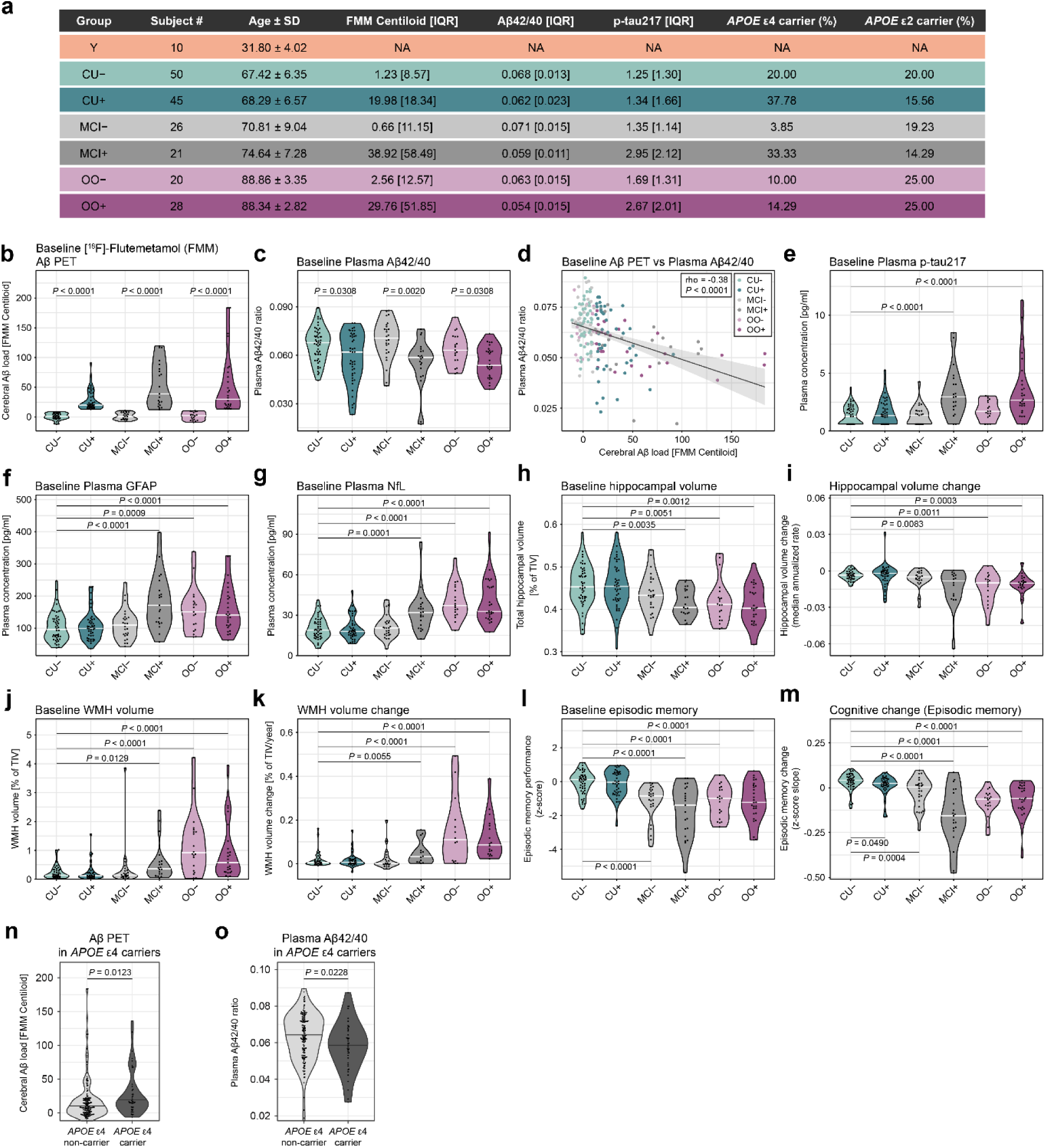
Baseline cohort characteristics and biomarker distributions across study participant subgroups. **a**. Tabular overview at a glance of the 200-participant study population, including young adults (Y), cognitively unimpaired older adults stratified by cerebral Aβ status (CU−, CU+), participants with mild cognitive impairment stratified by cerebral Aβ status (MCI−, MCI+), and oldest-old, 85+ years old participants without dementia stratified by cerebral Aβ status (OO−, OO+). Shown are number of subjects, baseline age (mean ± SD), baseline [18F]-flutemetamol (FMM) PET Aβ load as Centiloid (median [IQR]), plasma Aβ42/40 ratio (median [IQR]), plasma p-tau217 concentration in pg/ml (median [IQR]), and the percentage of *APOE* ε4 and *APOE* ε2 carriers. **b-m**. Baseline and longitudinal biomarker distributions across participant subgroups. Violin plots show baseline cerebral Aβ load measured by FMM Aβ PET (b), baseline plasma Aβ42/40 ratio (c), baseline plasma p-tau217 (e), baseline plasma GFAP (f), baseline plasma NfL (g), baseline hippocampal volume normalized to total intracranial volume (TIV) (h), longitudinal hippocampal volume change (i), baseline white matter hyperintensity (WMH) volume normalized to TIV (j), longitudinal WMH volume change (k), baseline episodic memory performance (l), and longitudinal episodic memory change (m). White lines indicate medians; points represent individual participants. Raw values are displayed. Statistical comparisons were performed using Dunn’s tests on sex-adjusted residuals, with *P* values adjusted using BH FDR. For cerebral Aβ load and plasma Aβ42/40 ratio (b,c), Aβ-positive and Aβ-negative participants were compared within each diagnostic group. For all other biomarkers (e-m), comparisons were performed against the CU− reference group. Association between baseline cerebral Aβ load and plasma Aβ42/40 ratio is shown in scatter plot (d). Grey line indicates a linear regression fit with 95% confidence interval. Correlation coefficients were calculated using Spearman’s rank correlation and are reported as ρ with corresponding *P* value; *P* values ≤ 0.1 are displayed. **n,o**. Violin plots showing baseline cerebral Aβ load (n) and plasma Aβ42/40 ratio (o) stratified by *APOE* ε4 carrier status across the pooled sample. Black lines indicate medians; points represent individual participants. Statistical comparisons were performed using Wilcoxon tests, with *P* values adjusted using BH FDR. Abbreviations: CU = cognitively unimpaired older adults; MCI = mild cognitive impairment; OO = oldest-old participants without dementia; FMM = [18F]-flutemetamol; IQR = interquartile range; GFAP = glial fibrillary acidic protein; NfL = neurofilament light chain; TIV = total intracranial volume; WMH = white matter hyperintensity; BH FDR = Benjamini-Hochberg false discovery rate approach.

**Extended Data Fig. 2:**
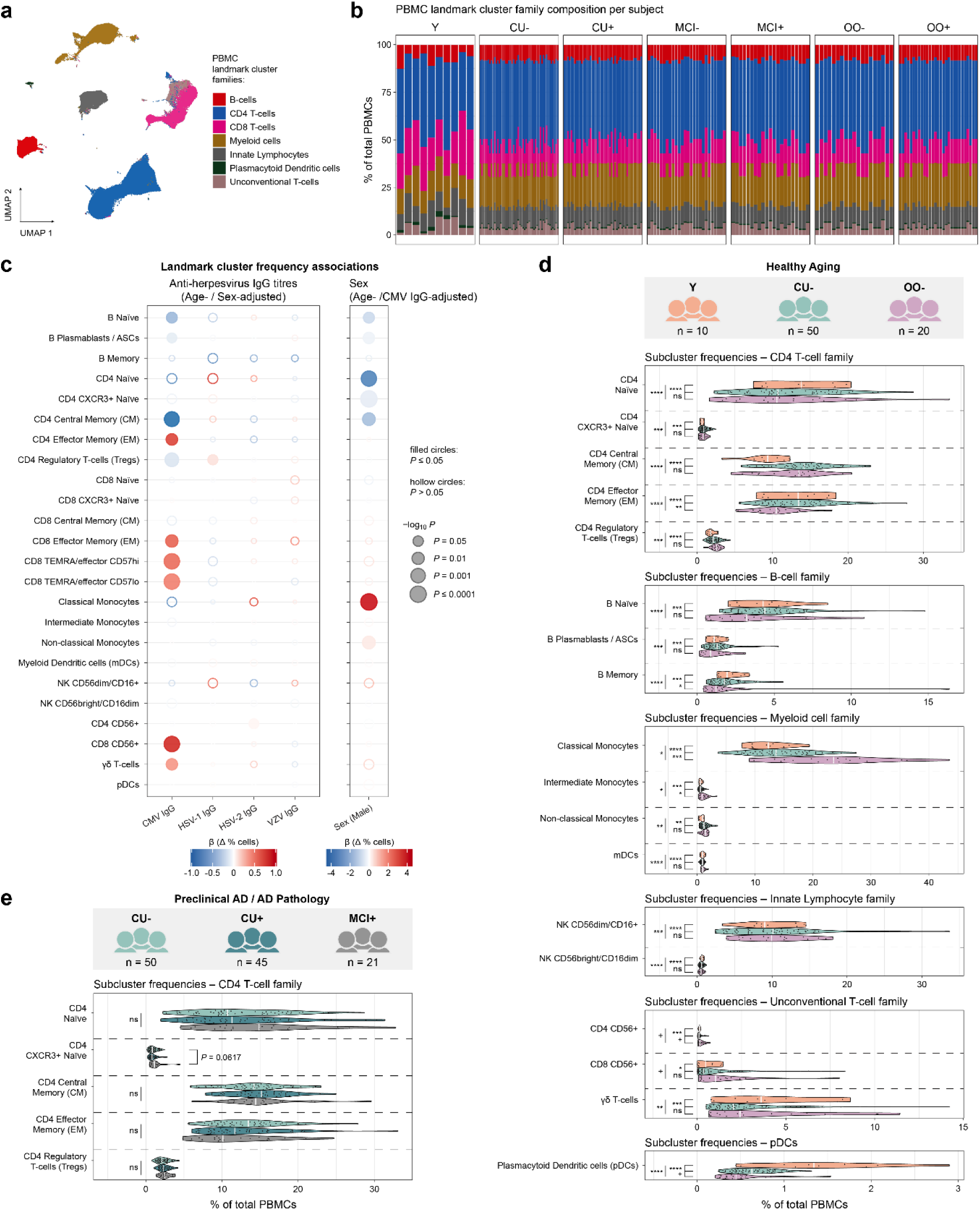
Global PBMC family structure, covariate associations and additional family-level immune cell cluster analyses. **a**. UMAP of total CD45+ PBMCs based on FlowSOM clustering. UMAP dimensionality reduction was computed on 10,000 cells; 5 million cells are displayed in final visualization. Colours indicate landmark cluster families. **b**. Stacked bar plots showing landmark cluster family composition per subject as percentage of total PBMCs, stratified by diagnostic and Aβ subgroup. **c**. Bubble plots summarizing linear models of landmark cluster frequency in relation to IgG titres against cytomegalovirus (CMV), herpes simplex virus 1 (HSV-1), herpes simplex virus 2 (HSV-2) and varicella zoster virus (VZV), and sex. Models for viral IgG titres were adjusted for age and sex; the model for sex was adjusted for age and CMV IgG titre. Bubble colour indicates the regression coefficient β and direction of effect, with red indicating higher and blue indicating lower landmark frequency. Bubble area is proportional to −log10(*P* value). Filled circles indicate *P* ≤ 0.05; hollow circles indicate *P* > 0.05. For viral IgG titres, β (Δ % cells) indicates the change in landmark cluster frequency, in percentage, per 1 unit increase in log-transformed IgG titre. For sex, β indicates the percentage difference in landmark cluster frequency in male compared with female participants. **d**. Healthy aging comparison including young adults (Y), PET Aβ-negative CU (CU−) and PET Aβ-negative OO (OO−), showing the remaining landmark cluster families not displayed in Fig. 2. Sideways violin plots show frequencies of CD4 T-cell, B-cell, myeloid cell, innate lymphocyte, unconventional T-cell and plasmacytoid dendritic cell family landmark nodes as percentage of total PBMCs. White lines indicate medians. Plots show raw values, whereas pairwise group comparisons were performed using Dunn’s test on hybrid values; CU− and OO− values were residualized for sex and CMV antibody titre, whereas Y values remained raw. *P* values were adjusted using BH FDR across all pairwise comparisons within each landmark node. BH FDR-adjusted *P* values ≤ 0.1 are displayed. ^+^ P < 0.1, * P < 0.05, ** P < 0.01, *** P < 0.001, **** P < 0.0001. **e**. Preclinical AD / AD pathology comparison including CU−, PET Aβ-positive CU (CU+) and PET Aβ-positive MCI (MCI+), showing CD4 T-cell family landmark-node frequencies as percentage of total PBMCs. White lines indicate medians. Pairwise group comparisons were performed using Dunn’s test, with all groups residualized for sex and CMV antibody titre. Testing was restricted to comparisons against the CU− reference group. *P* values were adjusted using BH FDR across the comparisons within each landmark node. FDR-adjusted *P* values ≤ 0.1 are displayed. Abbreviations: PBMCs = peripheral blood mononuclear cells; CU = cognitively unimpaired older adults; MCI = mild cognitive impairment; OO = oldest-old participants without dementia; UMAP = uniform manifold approximation and projection; BH FDR = Benjamini-Hochberg false discovery rate approach.

**Extended Data Fig. 3:**
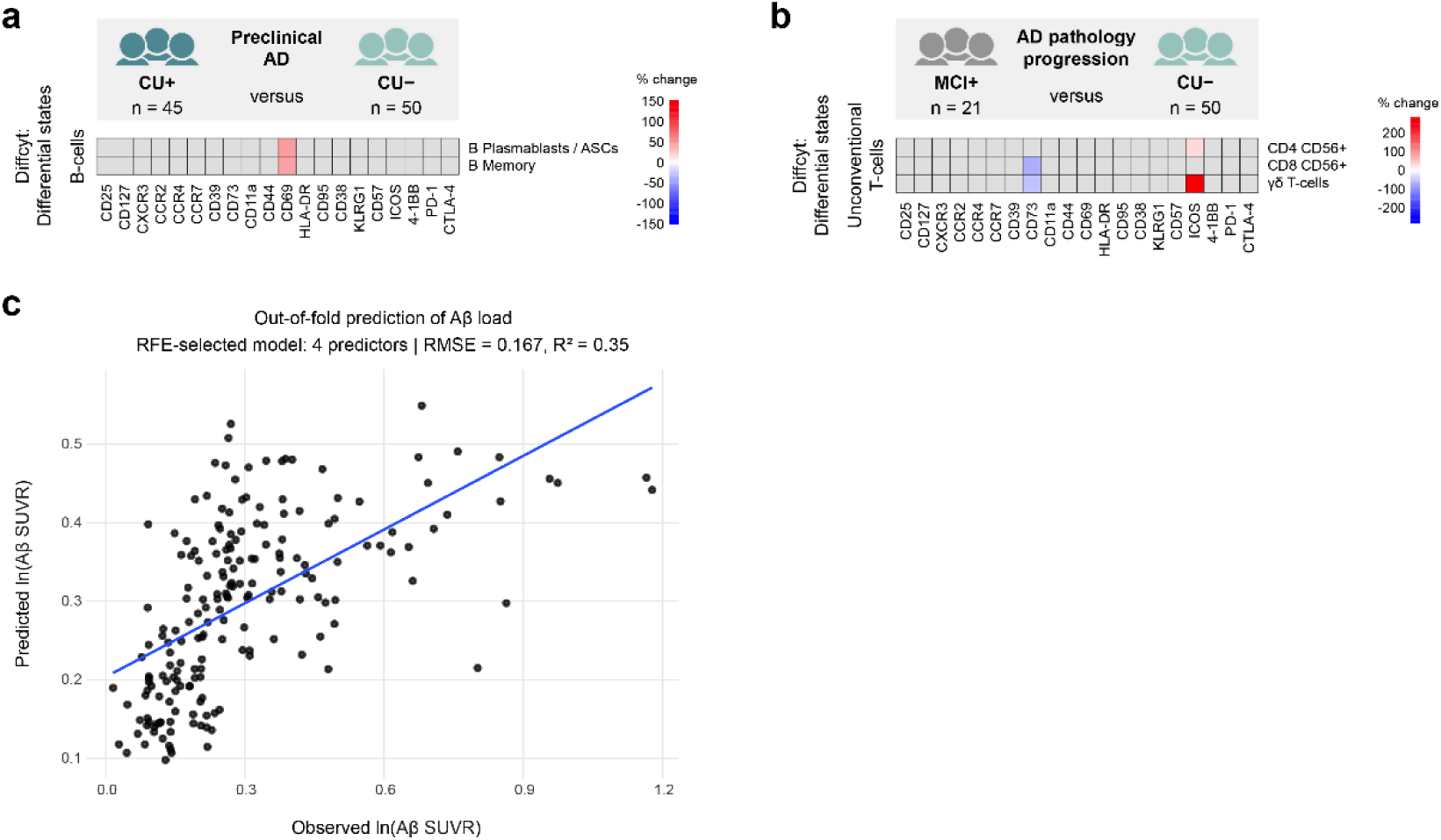
Additional Aβ pathology-associated immune cell states and cross-validated prediction of Aβ load. **a,b**. Additional Diffcyt differential state results from disease-relevant contrasts shown in Fig. 3. Heatmaps show significant cluster-marker features, including B-cell differential state hits in the ‘Preclinical AD’ contrast comparing PET Aβ-positive and Aβ-negative CU (CU+ versus CU−; a), and unconventional T-cell differential state hits in the ‘Pathology progression’ contrast comparing PET Aβ-positive MCI with PET Aβ-negative CU (MCI+ versus CU−; b). Analyses were adjusted for sex and CMV IgG. Colours indicate raw percentage change in marker expression. Grey tiles indicate features not passing BH FDR-adjusted threshold of *P* < 0.05; landmark clusters without significant hits are not shown. **c**. Cross-validated predicted versus observed cerebral Aβ load for the RF-RFE-selected immune marker model shown in Fig. 3. Each point represents one participant’s out-of-fold prediction from a random-forest model trained on the RFE-selected immune predictors. Blue line indicates a linear fit for visual guidance. Model performance was summarized across all out-of-fold predictions using RMSE and R², calculated as the squared correlation between observed and predicted ln(Aβ SUVR). Abbreviations: CU = cognitively unimpaired older adults; MCI = mild cognitive impairment; BH FDR = Benjamini-Hochberg false discovery rate approach; RF-RFE = random forest recursive feature elimination; RMSE = root mean squared error; SUVR = standardized uptake value ratio.

**Extended Data Fig. 4:**
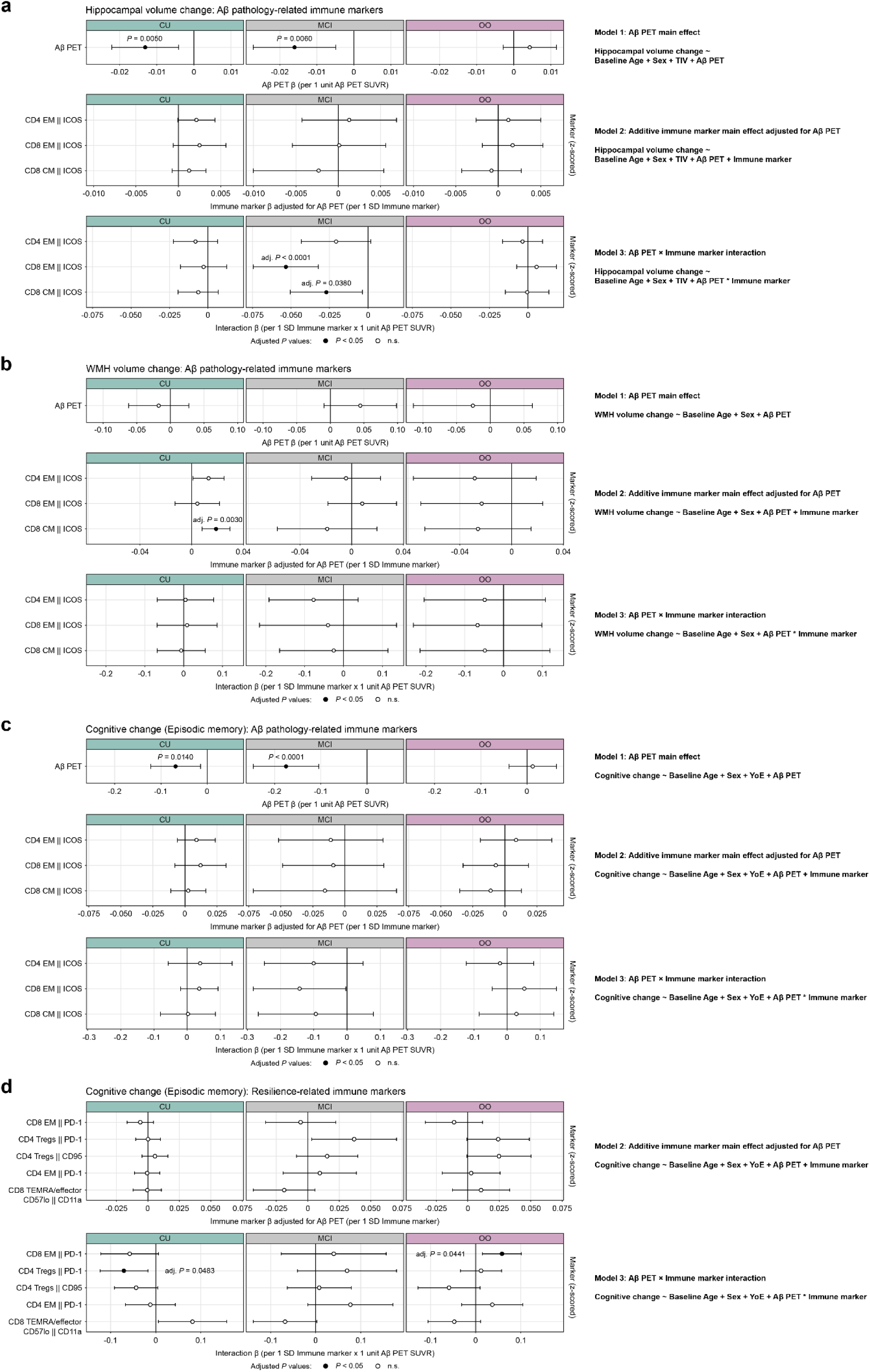
Linear model analyses of Aβ and immune-marker associations with longitudinal outcomes. **a-d**. Linear models testing associations of cerebral Aβ load, immune-marker expression and Aβ PET × immune-marker interactions with longitudinal outcome measures. Models were fitted separately within CU, MCI and OO. For each outcome, three model levels were assessed: Model 1 tested the Aβ PET main effect; Model 2 tested the additive immune-marker main effect adjusted for Aβ PET; and Model 3 tested the Aβ PET × immune-marker interaction. All models were adjusted for model-specific covariates. Model 1 is shown as one Aβ PET effect per diagnostic group; Models 2 and 3 are marker-specific. Forest plots show β estimates and 95% confidence intervals. For Model 1, β indicates the outcome change per 1-unit higher Aβ PET SUVR. For Model 2, β indicates the outcome change per 1 SD higher marker expression, adjusted for Aβ PET. For Model 3, β indicates the change in the Aβ PET outcome association per 1 SD higher marker expression. In Model 1, filled circles indicate *P* < 0.05. In Models 2 and 3, filled circles indicate BH FDR-adjusted *P* < 0.05 within each diagnostic group across all tested immune markers. **a**. Models of hippocampal volume change using Aβ pathology-related immune markers identified in Fig. 3. Models were adjusted for baseline age, sex and TIV. **b**. Models of WMH volume change using Aβ pathology-related immune markers identified in Fig. 3. Models were adjusted for baseline age and sex. **c**. Models of episodic memory change using Aβ pathology-related immune markers identified in Fig. 3. Models were adjusted for baseline age, sex and years of education. **d**. Episodic memory change models using resilience-related immune markers identified in Fig. 5. Models were adjusted for baseline age, sex and years of education. Abbreviations: CU = cognitively unimpaired older adults; MCI = mild cognitive impairment; OO = oldest-old participants without dementia; SUVR = standardized uptake value ratio; TIV = total intracranial volume; WMH = white matter hyperintensity; SD = standard deviation; BH FDR = Benjamini-Hochberg false discovery rate approach.

